# Structure-function analysis of enterovirus protease 2A in complex with its essential host factor SETD3

**DOI:** 10.1101/2022.06.22.497068

**Authors:** Christine E. Peters, Ursula Schulze-Gahmen, Manon Eckhardt, Gwendolyn M. Jang, Jiewei Xu, Ernst H. Pulido, Melanie Ott, Or Gozani, Kliment A. Verba, Ruth Hüttenhain, Jan E. Carette, Nevan J. Krogan

**Author notes:** Corresponding authors. Corresponding Authors K.A.V., R.H., J.E.C., N.J.K. Contributed equally (Christine E. Peters, Ursula Schulze-Gahmen).

## Abstract

Enteroviruses cause a number of medically relevant and widespread human diseases with no approved antiviral therapies currently available. Host-directed therapies present an enticing option for this diverse genus of viruses. We have previously identified the actin histidine methyltransferase SETD3 as a critical host factor physically interacting with the viral protease 2A. Here, we report the 3.5 Å cryo-EM structure of SETD3 interacting with coxsackievirus B3 2A at two distinct interfaces, including the substrate-binding surface within the SET domain. Structure-function analysis revealed that mutations of key residues in the SET domain resulted in severely reduced binding to 2A and complete protection from enteroviral infection. Our findings provide insight into the molecular basis of the SETD3-2A interaction and a framework for the rational design of host-directed therapeutics against enteroviruses.

## Introduction

Enteroviruses (EVs) comprise a genus of single-stranded RNA viruses of positive polarity whose members cause widespread human disease with large medical and economic impact^1^. They are the most common etiological agents of viral respiratory and neurological infections, usually causing mild symptoms but are also capable of causing severe pathology including asthma exacerbations, acute flaccid paralysis and myocarditis. There are currently no approved antiviral therapies available, partially due to a wide phenotypic diversity of enteroviruses^2^. Viruses rely on their host cells for replication, and have thus evolved strategies to modify the host cell environment to their advantage, often facilitated by virus-host protein-protein interactions. Host-directed therapy (HDT) aims to make use of this knowledge by interfering with host cell proteins that are critical to viral infection, replication or pathogenicity^3^. This strategy has the potential for broad-spectrum therapeutics by targeting cellular pathways and processes that are commonly exploited within virus families or genera. Furthermore, HDT may limit the emergence of drug resistance over traditional antivirals that target viral enzymes^4^, although this can depend on the particular host factor targeted and their role in the replication cycle^5^. Because of their diversity and high mutation rate^6^, EVs represent an intriguing target for HDT.

Systems biology approaches are especially valuable in the discovery of potential HDT targets for infectious diseases^7^. Through unbiased genetic screens, we have previously identified SETD3 as critically important for the replication of a broad range of enteroviruses including genetically diverse rhinoviruses, coxsackieviruses, poliovirus, and the non-polio enteroviruses EV-D68 and EV-A71, which are associated with severe neuropathogenesis^8^. SETD3 is a selective actin histidine methyltransferase that catalyzes mono-methylation of actin at histidine 73 (actin H73me) to regulate actin function^9, 10^. SETD3 belongs to a subclass of protein methyltransferases that in addition to containing the catalytic SET domain also have a Rubisco-substrate binding (RSB) domain that is present in the plant enzyme RLSMT (Rubisco large subunit methyltransferase). Human cells lacking SETD3 and mice with homozygous knockout alleles for *setd3* are both unable to support enteroviral RNA genome replication and hence are completely protected against viral pathogenesis. However, the methyltransferase activity of SETD3 is not required for its role in viral replication, indicating that SETD3 has a function in EV replication independent of actin methylation^8^.

Affinity purification mass spectrometry (AP-MS) has proven a powerful tool for identifying protein-protein interactions (PPIs) underlying diseases, including for several viral infections^11–13^. Similarly, our previous AP-MS studies revealed that enteroviruses co-opt SETD3 through a PPI with the viral 2A non-structural protein^8^. Enterovirus 2A is a protease that processes the viral polyprotein, and also cleaves a variety of host proteins^14^. This multi-functional protein is essential for RNA replication through protease-dependent^14, 15^ and independent^16, 17^ mechanisms. While the exact role of SETD3 during this process is incompletely understood, critical non-catalytic functions mediated through PPIs have been described for other methyltransferases^18, 19^. We thus propose that SETD3 is a master regulator of enterovirus pathogenesis through its interaction with 2A.

SETD3 is a promising target for HDT development as its role as a pan-enterovirus host factor would allow for the targeting of multiple different EV infections. Notably, SETD3 knock-out mice were viable^8, 10^, suggesting that short-term inhibition might have limited toxicity. To obtain a detailed understanding of the molecular interaction, we determined the cryo-EM structure of SETD3 in complex with coxsackievirus B3 (CV-B3) 2A at 3.5 Å resolution, revealing extended interactions between 2A and SETD3 residues, including some located in the substrate binding region of SETD3^10^. Analysis of the complex structure revealed two distinct 2A interaction interfaces on SETD3, one in the SET domain that overlaps with the SETD3 actin binding site, and one in the RSB domain. Mutations in the RSB domain interface had only minor effects on 2A binding and no effects on viral replication while mutations of key contact residues in the SET interface severely reduced binding affinity to 2A and completely abrogated viral replication. Interestingly, the binding site of SETD3’s canonical substrate, actin, and 2A on SETD3 partially overlap and show structural similarities in forming a short intermolecular β-sheet in both complexes. In addition, binding of SETD3 to 2A is inhibited in the presence of actin substrate peptide. These key structural insights into the interactions between the cellular methyltransferase SETD3 and enteroviral 2A may provide a framework for the rational design of therapeutics against this class of viruses of strong medical importance.

## Results

### CV-B3 2A and SETD3 form a stable complex

We have previously shown that the interaction of SETD3 with the enterovirus protease 2A is essential for enterovirus replication^8^. However, whether SETD3 binds directly with 2A or is part of a larger protein complex is unknown. To investigate this, we expressed SETD3 in human cells as a fusion protein with a FLAG affinity tag and affinity-purified SETD3 in the presence and absence of CV-B3 2A. Since the expression of catalytically active 2A protease cleaves multiple host cellular proteins leading to cell toxicity^20^, all 2A constructs used in this report are catalytically inactive mutants, unless otherwise stated. Specifically, we used a CV-B3 2A mutant that encodes a substitution of the active-site cysteine (2A_C107A_). Interacting host proteins that associate with SETD3 were identified in an unbiased manner using quantitative mass spectrometry (MS). High-confidence interactors were identified using the SAINTexpress (Significance Analysis of INTeractome) algorithm^21, 22^ and their abundances were compared in the presence and absence of 2A using MSstats^23^. SETD3 interactions with host proteins did not change upon introduction of 2A, suggesting that 2A binding to SETD3 does not recruit any additional host factors (**Fig. 1A** and **Table S1**). To further probe this hypothesis and determine if other host proteins associate with the SETD3-2A complex, we performed a sequential affinity purification as previously described^24, 25^, followed by quantitative MS (**Fig. 1B**). We confirmed a complex between SETD3 and 2A with no other stable protein components (**Fig. 1C** and **Table S2**). Not only were SETD3 and 2A the only proteins detected with statistical significance, peptides spanning the vast majority of their sequence were recovered. Furthermore, we compared the estimated abundances of SETD3 and 2A from the sequential affinity purifications, which strongly suggest a 1:1 stoichiometry of the SETD3-2A complex (**Fig. S1A**).

**Figure 1.**
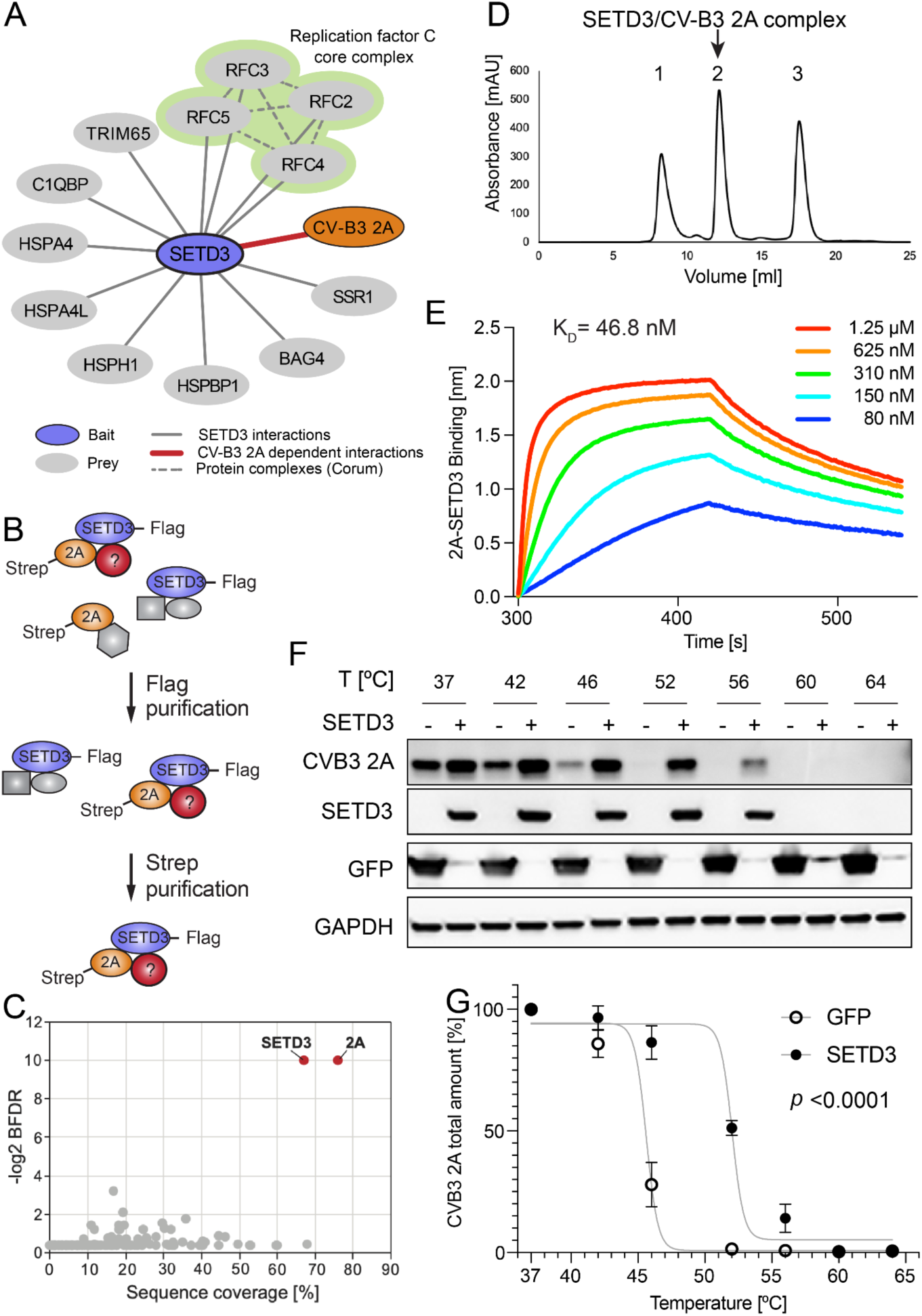
CV-B3 2A and SETD3 form a stable complex. **A**. High-confidence SETD3 interacting host proteins in presence and absence of CV-B3 2A with a BFDR <0.05 generated using AP-MS. See also **Table S1. B**. Schematic of sequential FLAG-Strep AP (double-AP) experimental flow. **C**. Dot plot representation of proteins identified in double-AP experiment. Sequence coverage of detected proteins versus SAINTexpress BFDR is shown. Proteins passing the significance threshold of BFDR <0.05 are highlighted in red. See also **Table S2**. **D**. Purification of reconstituted SETD3-2A complex. Size exclusion column elution profile after purification and complex reconstitution is shown. Aggregated material (peak 1), complex of 2A-protease with SETD3 (peak 2), and monomeric 2A (peak 3). Purified complex from peak 2 was used for structure determination by cryo-EM. See also **Fig. S1**. **E**. Binding of purified SETD3 to 2A protein as measured by biolayer interferometry. See also **Fig. S1+5**. Binding curves for varying input concentrations (different colors indicated in legend to individual plots) of SETD3. Equilibrium constant K_D_ as calculated by the integrated software of the Octet Red384 instrument is shown. **F.** Stability assay for 2A protein. Representative Western blot analysis of CETSA for 293FT cells co-transfected with CV-B3 2A and SETD3-Flag or GFP-Flag. **G.** Quantification of protein stability. CETSA melting curves of CV-B3 2A upon co-transfection with SETD3-Flag or GFP-Flag are shown for three biological replicates. Samples incubated at 42°C were only taken in duplicate. Western blot band intensities of 2A incubated at 37°C were normalized to gapdh and set as 100%. 2A protein intensities normalized to gapdh were related to the signal at 37°C. Relative 2A intensities were plotted against corresponding incubation temperatures and a non-linear regression was applied. Data represent the mean and standard deviation. *P* values were calculated using GraphPad Prism.

To further determine the molecular details of this interaction, we recombinantly expressed SETD3 and CV-B3 2A individually in *E. coli,* followed by reconstitution of the complex from the individually purified subunits. SETD3 and 2A formed a stable complex (**Fig. 1D**, peak 2, **Fig. S1B**) with a K_D_ of 47 nM as determined by biolayer interferometry (BLI) (**Fig. 1E**). Next, we examined the effect of SETD3 binding on 2A stability by a cellular thermal shift assay (CETSA). This allows for the investigation of protein complexes in intact mammalian cells based on the premise that bonafide interaction partners increase protein stability upon heat denaturation^26, 27^. Co-expression of 2A with SETD3 resulted in a marked increase of 2A protein levels compared to co-expression with a control protein (GFP). This suggests that complex formation with SETD3 influences 2A stability and prevents 2A degradation. Following heating of the cells at various temperatures, the amount of soluble 2A protein gradually decreased in cells co-expressing GFP indicative of protein aggregation due to heat denaturation. Strikingly, in the presence of SETD3, 2A remained soluble at higher temperatures (**Fig. 1F**) resulting in a shifted melting curve (**Fig. 1G**). Our results indicate that the binding interaction with SETD3 increases 2A cellular stability, which could have functional consequences on the multifunctional protein 2A’s role in viral replication. Taken together, these results indicate that SETD3 and 2A form a stable and stoichiometric (1:1) complex, which prompted us to perform detailed functional and structural studies on this virus-host interaction.

### Structure of SETD3 in complex with CV-B3 2A

We determined the SETD3-CV-B3 2A complex structure to 3.5 Å resolution by cryo electron microscopy (cryo-EM) and observed clear density for residues 22-495 of SETD3 and 4-141 of 2A protease (**Fig. 2A**, **Figs. S2-S4** and **Table S3**). Similar to previous crystal structures of SETD3, residues 496-594 were not visible in the cryo-EM density map. CV-B3 2A binds in the pocket formed by the SET domain and the RSB domain of SETD3, forming two separate interaction interfaces with SETD3. The interaction surface areas with the SET domain and the RSB domain including the linker are 747 Å^2^ and 460 Å^2^, respectively. Contacts in the RSB interface are more solvent accessible due to the smaller interaction surface (**Fig. 2B**), including van der Waals contacts and hydrophobic interactions, but no hydrogen bonds. The linker region connecting the SET domain and the RSB domain contributes two hydrogen bonds/salt bridge contacts between SETD3 R336 and 2A E74 (**Table S4**). In contrast, the interaction surface with the SET domain is much larger and includes a small hydrophobic core area centered on interactions between SETD3 residues Q257 and P259, and 2A residue V59. This core area is surrounded by more hydrophilic interactions, including a few solvent exposed side-chain-hydrogen bonds with 2A (**Fig. 2B+C** and **Table S4**). Additionally, a pair of main-chain main-chain hydrogen bonds between SETD3 Y288 and 2A H68 form part of an anti-parallel intermolecular β-sheet structure between the two proteins (**Fig. 2C**). Notably, of the 2A residues in the SET domain binding interface (SETD3 residues 94-314) observed in our cryo-EM structure, the majority overlap with 2A residues identified as critical for enterovirus replication using a mammalian two-hybrid system (bold residues in **Table S4**)^8^.

**Figure 2.**
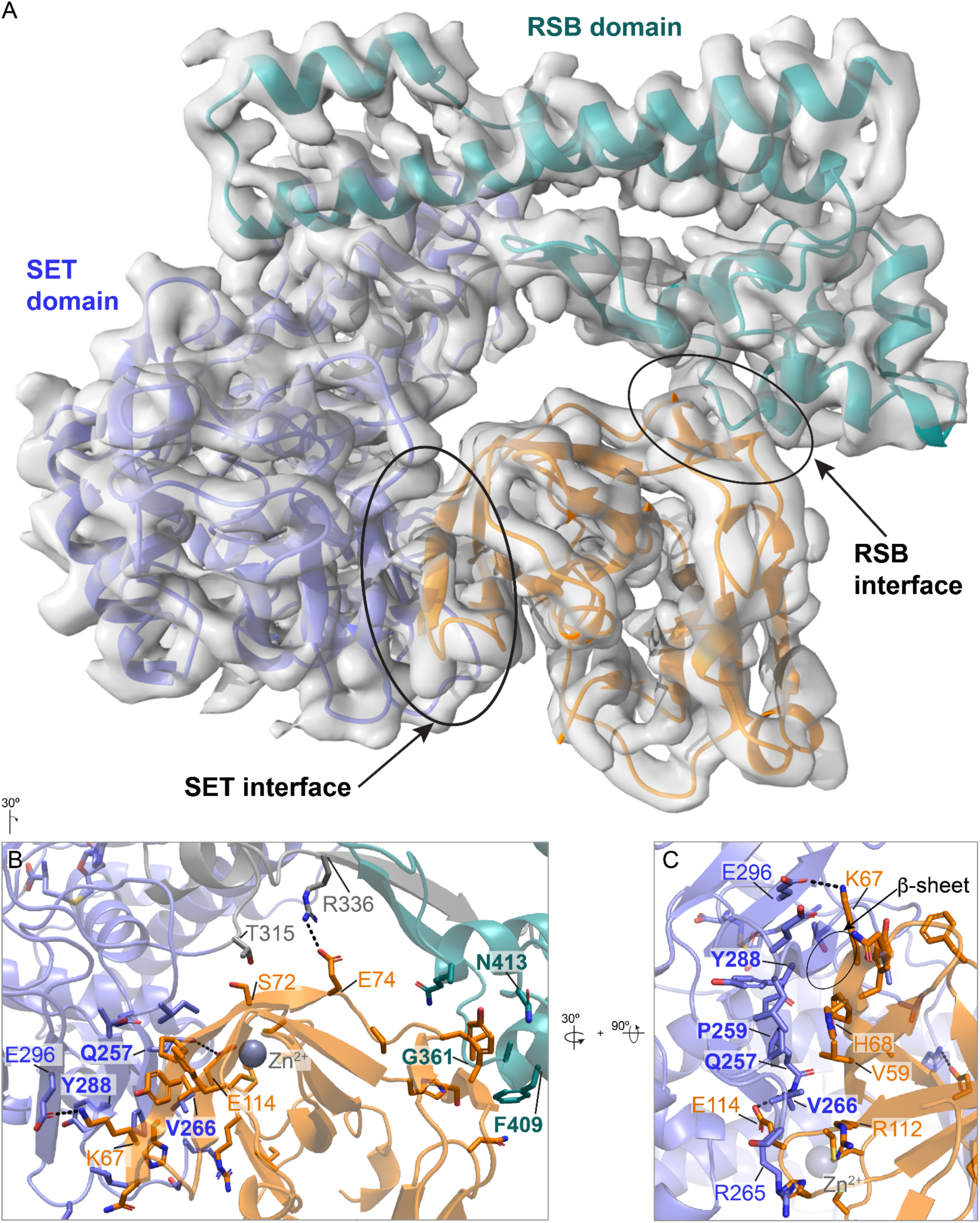
Structure of CV-B3 2A and SETD3 in complex. **A**. Cryo-EM density and ribbon representation of SETD3-2A complex. The SET domain of SETD3 is shown in purple, the linker in gray, the RSB domain in cyan, and CV-B3 2A protease in orange. See also **Table S3, Table S4** and **Figs. S2-S4**. **B**-**C**. Expanded views of SET interface and RSB interface interactions, with important residues highlighted. Indicated contact residues of SETD3 and 2A are shown as stick representation. SETD3 residues selected for mutational studies (see Fig. 3 and onwards) are labeled in bold font.

### The SET interface is critical for 2A binding

We have previously shown that mutations in 2A residues that abolish binding to SETD3 inhibit viral replication when introduced into the viral genome^8^. However, we cannot exclude that those mutations disrupted another function of 2A in addition to binding to SETD3, or that they are generally destabilizing the structure of 2A protease. Therefore, to test our structural observations, we focus on functionally characterizing key interaction-defining residues from the two interaction interfaces on SETD3 (**Fig. 3A**). Substitutions were made in SETD3 residues that form more than 10 contacts with 2A (Q257A, V266D, Y288P, F409A, N413A) as well as G361R, which is in the RSB domain interface. A glycine to arginine mutation in SETD3 residue 361 is expected to disrupt the SETD3-2A interaction by introducing steric clashes (**Fig. 3B**). In general, the specific amino acid replacements were chosen to maximize the effect of the mutation on the respective residue’s interactions.

**Figure 3.**
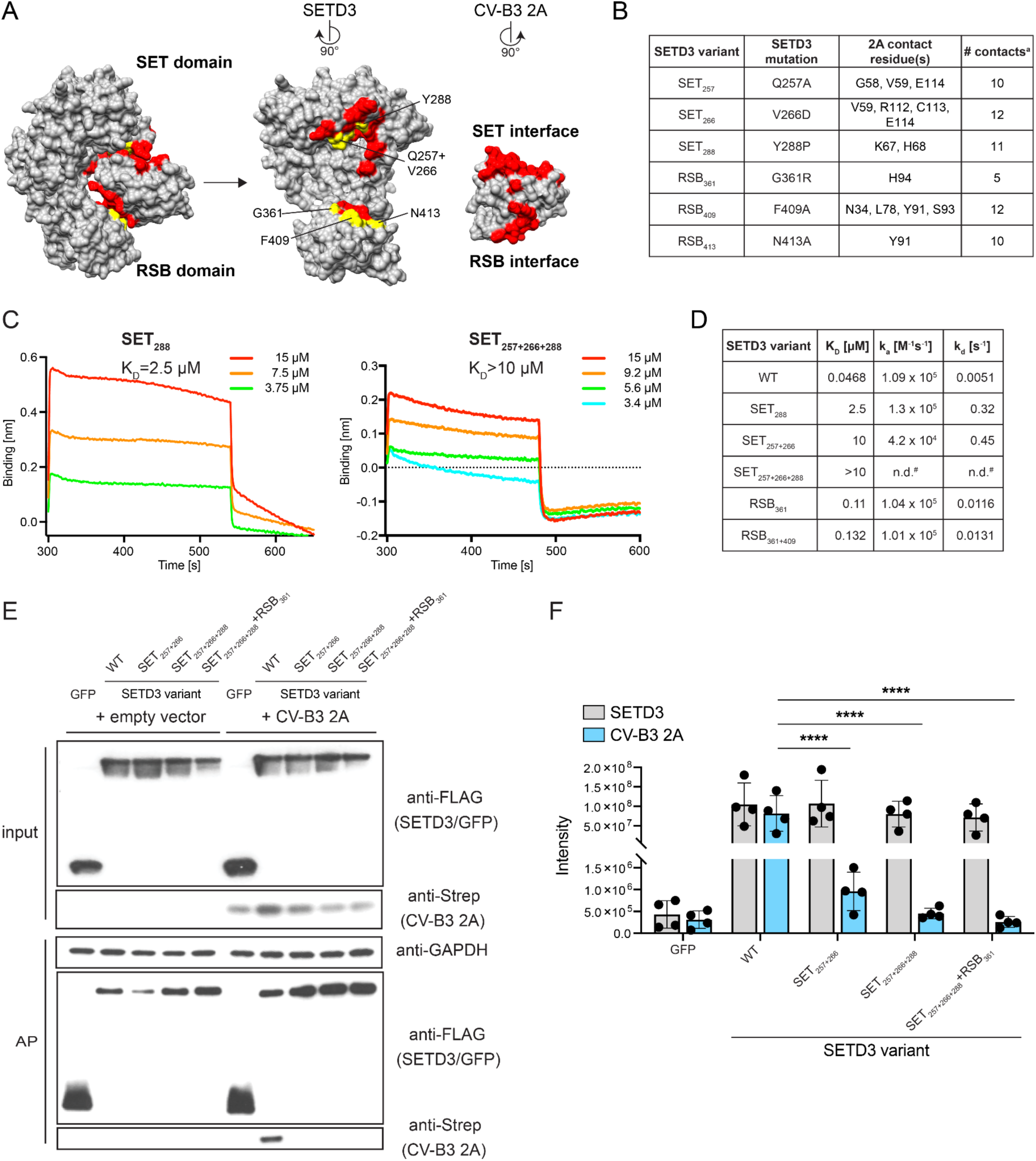
The SET interface is critical for 2A binding. **A**. Surface view of the binding interface and the selected mutants for functional validation. Residues making up the interaction interface on SETD3 and 2A are labeled in red. Residues on SETD3 mutated for further analysis are indicated in yellow. See also **Table S4**. **B**. Characteristics of the residues selected for mutations. ^a^Contact defined as <4.1Å. See also **Table S4**. **C-D**. *In vitro* biolayer interferometry binding assay results for purified SETD3 variants to 2A protein. See also **Fig. S5**. **C**. Binding curves for varying input concentrations (different colors indicated in legend to individual plots) of selected SETD3 variants. Equilibrium constants K_D_ as calculated by the integrated software of the Octet Red384 instrument are shown. **D**. Summary table of *in vitro* binding assay. Rate constants and equilibrium constants were determined using the integrated software of the Octet Red384 instrument. n.d. = not determined; ^#^Binding curves showed reduced binding compared to SET_257+266_ mutant. Exact values could not be calculated for this low affinity binding. **E**. Co-AP-WB analysis of SETD3 variants binding to CV-B3 2A. Input and eluates after Flag-affinity purification (AP) are shown. **F**. Quantitative MS analysis of SETD3 variants binding to co-transfected CV-B3 2A. Protein intensities for SETD3 and 2A, as determined by label-free quantitative MS, are shown for four technical replicates. Bars and error bars represent mean and standard deviation, respectively. Statistical models implemented in MSstats were used to calculate *P*-values, which were adjusted with Benjamini Hochberg. \*\*\*\**P* ≤ 0.0001.

To measure the effect of SETD3 mutants on complex formation with 2A, we determined *in vitro* binding equilibrium and rate constants using BLI (**Fig. 3C+D, Fig. S5A**). Compared to the high affinity binding between WT SETD3 and 2A (K_D_ = 47 nM), a single mutation of Y288P in the SET domain (SET_288_) reduces binding affinity 53-fold (K_D_ = 2.5 µM) and a double mutation in the same domain, SET_257+266_, 213-fold (K_D_ = 10 µM). BLI measurements for the triple mutant in the SET interface, SET_257+266+288_, showed even lower affinity binding events that could not reliably be used to determine a K_D_ value. In contrast, the single and double mutants in the RSB domain of SETD3, RSB_G361_ and RSB_G361+F409_, showed smaller reduction in binding affinities with a calculated K_D_ of 110 nM and 132 nM, respectively. The integrity of all SETD3 mutants was confirmed by comparing size exclusion chromatography elution profiles of the final purification step. All mutants eluted at very similar elution volumes, indicating no major changes in the protein structure (**Fig. S5B)**.

We next sought to test SETD3-2A complex formation in the context of cells, focusing on mutants of the SET interface that had substantially decreased *in vitro* binding. To this end, we co-expressed FLAG-tagged SETD3 mutants and Strep-tagged CV-B3 2A in 293T cells and performed a FLAG affinity purification (AP). As expected, 2A was readily observed in the eluate of the SETD3 WT AP upon Western blotting. In contrast to this, no detectable 2A copurified with SET interface double and triple mutants (SET_257+266_ and SET_257+266+288_), as well as a quadruple mutant including an additional mutation in the RSB interface (SET_257+266+288_+RSB_361_) (**Fig. 3E**). To quantify these results, the eluate was analyzed using quantitative MS. While protein intensities showed that all SETD3 variants were enriched equally, 2A enrichment in SET interface mutant affinity purifications was significantly decreased, suggesting a loss of robust protein-protein binding (**Fig. 3F**). In summary, mutations in the SET domain of SETD3 had a detrimental effect on CV-B3 2A binding, especially when multiple residues were affected. The critical role of SETD3 on viral replication could potentially involve regulation of 2A proteolytic activity. To evaluate the effect of SETD3 binding on protease activity, we measured the specific activity of 2A towards a peptide covering the cis-cleavage site between VP1 and 2A (**Fig. S5C**). Upon binding with WT SETD3, we found a 1.5 fold increase of 2A activity with k_cat_/K_m_= 1.5 mM^-1^ min^-1^ for 2A alone and 2.28 mM^-1^ min^-1^ for 2A-SETD3 complex, respectively. In contrast, the presence of the non-binding triple mutant SETD3_257+266+288_ was not able to increase 2A activity.

### 2A binding partially overlaps with the substrate binding site at the SET interface

The SET domain we identified as critical for SETD3-2A complex formation has previously been shown to be involved in substrate binding of SETD3’s canonical substrate, actin^10^. Specifically, a synthetic actin-peptide substrate composed of beta-actin residues 66-80 binds on the surface of the SET domain with actin H73 and G74 adopting a turn structure that positions the H73 side-chain in the SETD3 active site to allow for methyl transfer^10^. Importantly, the substrate binding site on SETD3 is sterically removed from its active site pocket (formed by conserved residues W274, Y313 and R316), as well as residues important for binding of the methyl donor S-adenosyl-methionine (conserved residues N278 and H279). To test if 2A interacts with SETD3 similarly to its substrate, actin, we superimposed the structure of the SETD3-actin peptide complex (PDB ID 6mbk) onto our cryo-EM structure of the 2A-SETD3 complex (**Fig. 4A**). The SETD3 molecules superimposed with a root mean square deviation (RMSD) of 1.08 Å for Cα-atoms. Although we do not observe any overlap of actin and 2A binding in the SETD3 active site, actin peptide residues 66-72 and 2A partially overlap in the substrate binding region, forming intermolecular β-sheets with SETD3 involving SETD3 residue Y288. The backbone amide and carbonyl oxygen of SETD3 Y288 form hydrogen bonds with H68 carbonyl and amide of 2A. In the actin complex, the SETD3 Y288 amide forms a hydrogen bond with the Y69 carbonyl in actin, and a second backbone hydrogen bond is formed between the SETD3 T286 carbonyl oxygen and the I72 amide in the actin peptide (**Fig. 4A**, inset). In both complexes, the intermolecular hydrogen bonds are stabilizing the complex between SETD3 and its ligand and are leading to a similar side-chain orientation for 2A H68 and actin Y69, suitable for hydrophobic interactions with SETD3 P259 and L290. Replacement of tyrosine with proline in the SET_288_ mutant prevents backbone hydrogen bonds with the amide nitrogen and likely causes a local structural change around residue 288 in SETD3, affecting 2A binding.

**Figure 4.**
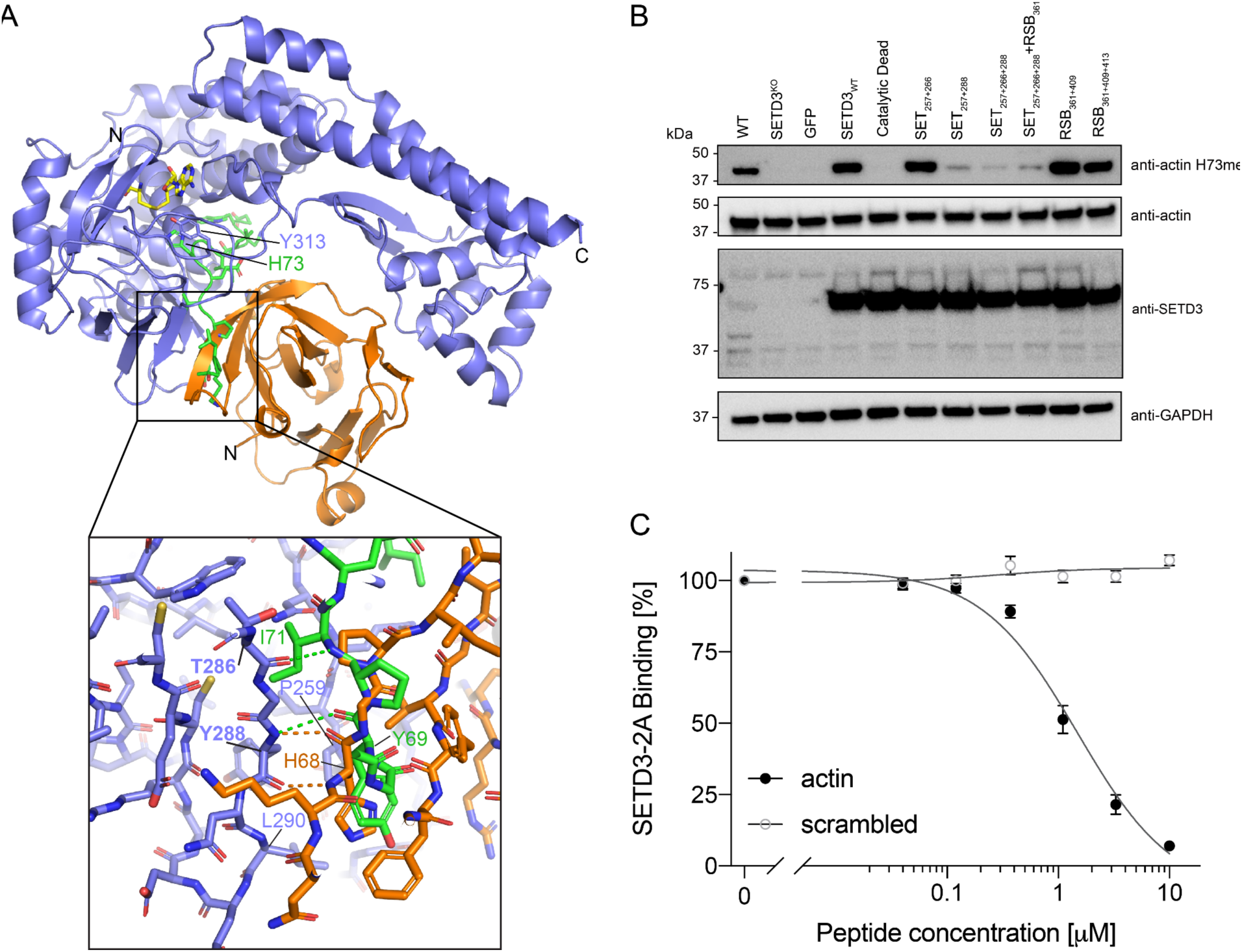
2A binding partially overlaps with the substrate binding site at the SET interface. **A**. Superposition of SETD3-2A cryo-EM structure and SETD3-actin complex (PDB ID 6mbk) with SETD3 from actin complex not shown. SETD3-2A ribbon model colors as in Fig. 2, β-actin bound to SETD3 is colored in green. S-adenosyl-l-homocysteine (SAH) in yellow, actin substrate residue H73, and SETD3 Y313 are shown as stick models, located in the active site of SETD3. Inset shows details of the intermolecular β-sheet structure observed between CV-B3 2A protease and SETD3 and between β-actin and SETD3, with important contact residues and hydrogen bonds indicated in the respective colors. **B**. Western blot analysis of actin methylation activity of WT, SETD3^KO^, and SETD3^KO^ H1-Hela cells stably complemented with the indicated SETD3 variants. Expression of SETD3 mutants is shown by Western blot. **C**. Impact of actin peptide on SETD3-2A binding in biolayer interferometry assay. Maximum SETD3 binding (310 nM) is plotted against peptide concentration.

Since SETD3 Y288 is involved in complex formation with both 2A and actin, this raises the possibility that the SET_288_ mutation will also impair the catalytic activity of SETD3. To test this, we compared the actin methylation status in cells in the presence of SETD3 WT or the structure-derived SETD3 mutants. We stably expressed the SETD3 constructs in a H1-HeLa^+CDHR3^ isogenic knock-out cell line of SETD3 (SETD3^KO^, **Fig. S6**) under a high expression promoter. As a control we used a previously described catalytic dead mutant of SETD3^8^ which contains five mutations in conserved residues that participate in forming the active-site pocket for actin methylation (W274, Y313 and R316) and conserved residues important for *S*-adenosyl-methionine (the methyl donor for methyltransferases) binding (N278 and H279). As expected, expression of SETD3 WT in SETD3^KO^ cells restored methylated actin levels to levels observed in wild-type cells, while the catalytic dead mutant failed to do so (**Fig. 4B**). Full actin methylation activity was detected for SETD3 containing mutations in the RSB interface and for the SET_257+266_ construct. As predicted, in cells expressing SETD3 constructs containing the Y288P mutation, strongly reduced methylated actin levels were observed suggesting that this mutation leads to partially inhibited methylation activity. This was not due to differences in SETD3 mutant protein expression as all mutants were expressed to higher levels than endogenous SETD3 in wild-type cells as confirmed by Western blot analysis (**Fig. 4B**).

The structural analogies between 2A and actin-peptide binding to SETD3 suggest that SETD3 cannot interact with its substrate actin while in complex with 2A. To test this hypothesis, we measured SETD3 binding to 2A in the presence of increasing amounts of actin peptide 66-81 with His73 replaced by leucine^28^. By keeping the protein concentrations of 2A and SETD3 constant while increasing the concentration of inhibitor peptide, we were able to measure reduced levels of maximum binding response with increasing peptide concentrations (**Fig. 4C**) in BLI experiments. Depending on SETD3 concentrations the IC50 values for inhibition of 2A binding by actin peptide ranged from 1.5 µM for 310 nM SETD3 to 3.4 µM for 625 nM SETD3, confirming a competition between actin and 2A binding to SETD3. Collectively, our results show that 2A binding to SETD3 uses a similar interface as its canonical substrate, actin, and key 2A interaction residues on SETD3 are also important for actin methylation. The actin peptide competes with 2A for binding to SETD3 *in vitro*, confirming our expectations that 2A and actin cannot bind simultaneously to SETD3.

### 2A binding at the SET interface is required for viral infection

We next tested which SETD3 residues in the 2A binding interface are critical for viral infection. To this end, we stably expressed the structure-derived SETD3 mutants in SETD3^KO^ cells under a minimal promoter. We then performed synchronized infection assays using an infectious CV-B3 virus expressing luciferase (CV-B3-RLuc) (**Fig. 5A**). For this, H1-Hela^+CDHR3^ cells, SETD3^KO^ cells, and SETD3^KO^ complemented cells were incubated in the presence of virus on ice for 1 h, and then at 37°C for 10 min to allow for synchronous viral entry. Cells were then either harvested to take a 0 h time point to detect background luminescence or cultured for an additional 8 h. This time point was chosen to capture virus replication in the exponential phase. We validated that all SETD3 constructs were expressed under the minimal promoter by Western blot analysis (**Fig. 5B**). Consistent with previous results, SETD3^KO^ cells showed a large decrease in viral replication compared to infection of WT cells (**Fig. 5C**). Complementation with either a SETD3 WT or a catalytic dead SETD3 restored viral infection similar to WT levels. Interestingly, addition of SETD3 containing mutations in the RSB interface were still able to restore viral infection to WT levels, suggesting that this interface has an inconsequential contribution to the binding interaction with 2A. While SET_257+266_ showed a large decrease in binding affinity to 2A (see **Fig. 2D-F**), complementation with this construct was still able to partially restore viral infection. Strikingly, the combination of three mutations in the SET interface (SET_257+266+288_) that led to a complete loss of binding, failed to restore viral infection.

**Figure 5.**
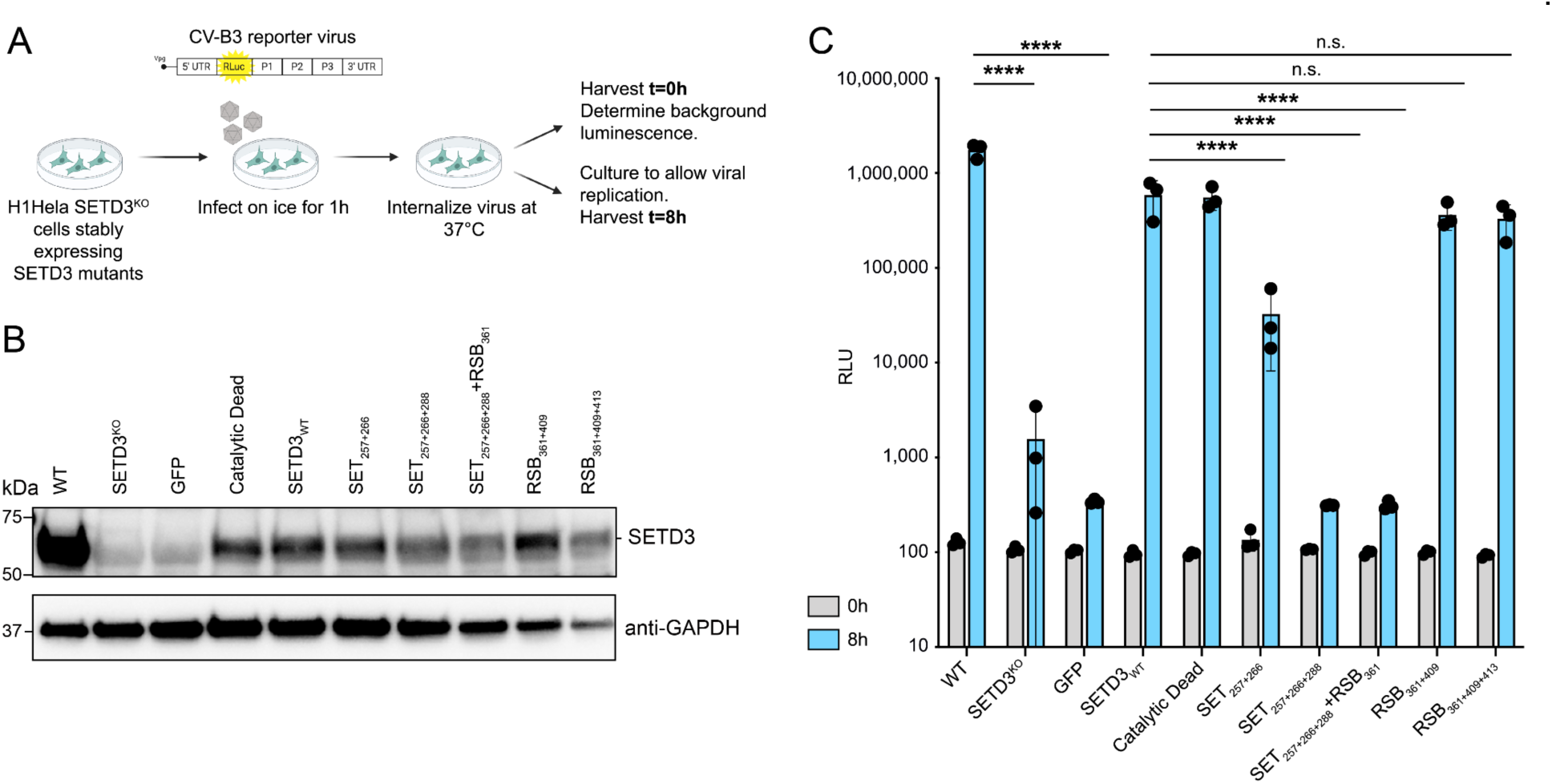
2A binding at the SET interface is required for viral infection. See also **Fig. S7. A**. Schematic of synchronized luciferase-expressing CV-B3 (MOI 1) infection. **B**. Western blot analysis of H1-Hela WT, SETD3^KO^, or SETD3^KO^ cells complemented with SETD3 variants under a minimal promoter are shown. **C**. Infection of WT, SETD3^KO^, or SETD3^KO^ cells complemented with SETD3 structure derived mutants. Statistics were performed on 8h timepoints. *P-*values were determined by two-way ANOVA (Holm–Sidak corrected) on log-transformed data. RLU = relative light units, n.s. = not significant, \*\*\*\**P* ≤ 0.0001.

Similar results were obtained with SETD3 mutant complementation using a promoter that drives higher levels of expression (CMV) (**Fig. S7**). Here, a 6 h time point was chosen, due to the faster replication kinetics with a higher expression of SETD3. Despite being overexpressed to much higher levels than endogenous SETD3 (compare **Fig. 4B**), the triple mutant SET_257+266+288_ was still unable to support viral infection. One difference seen is a full restoration of viral replication upon complementation with SET_257+266_ upon higher expression. While this mutant has a reduced binding affinity to 2A, it is likely that the high level of expression under the CMV promoter compensates for this reduction in binding. All together these data indicate that the SET interface is critical for SETD3 binding to 2A, thus supporting viral infection.

## Discussion

Here we report a cryo-EM structure of CV-B3 2A in complex with SETD3, revealing 2A bound in the cavity formed by the SET and RSB-domains of SETD3. We found that 2A interacts with SETD3 at two distinct interfaces, one present in the SET domain and the other in the RSB domain. Structure-function analysis revealed that the SET interface is critical for 2A binding and that 2A and actin peptide (66-81) bind competitively to the SET domain. Mutation of key SET domain residues resulted in severe loss of 2A binding and a complete loss of enteroviral infection. Our findings demonstrate the centrality of this virus-host protein-protein interaction in the life cycle of this enterovirus and uncover a molecular mechanism for how viruses have evolved to co-opt their host.

Affinity purification of SETD3 expressed in human cells in addition to binding experiments using purified SETD3 and 2A suggest that SETD3 promotes viral replication via direct binding and without recruiting additional host factors. This is in contrast to another well-established enterovirus-host interaction, where the Golgi-localized viral protein 3A interacts directly with the adaptor protein ACBD3, which recruits PI4KB required for the formation of the membranous vesicles that support viral replication^29–32^. The strong interaction of SETD3 with viral 2A is evidenced by its low dissociation constant (K_D_ = 47 nM). This is comparable to the binding affinity of SETD3 with its actin substrate (K_D_ = 170 nM), experimentally determined with a peptide containing the target histidine^33^. Mutations in the RSB interaction interface only modestly affected 2A binding whereas mutations in the SET interface severely affected binding. This interface partially overlapped with the core domain in SETD3 that binds actin with residue 288 involved in actin interactions as well as 2A interactions. Indeed, mutation of this residue not only affected binding with 2A but also reduced, but not ablated, the ability of SETD3 to methylate actin in H1-HeLa cells. Our data suggest that enteroviruses have evolved to engage SETD3 in a manner mimicking the way it interacts with its physiological substrate. Because 2A is not methylated by SETD3 in an *in vitro* assay^8^, this viral mimicry does not involve protein methylation. This is further supported by the observation that mutations in the catalytic pocket of SETD3 that do not interfere with the 2A binding domain, do not affect viral replication^8^. Recently, it has been shown that protein methyltransferases can regulate cellular pathways in a methyltransferase-independent manner primarily through facilitating protein-protein interactions^18, 19^. By structurally and functionally characterizing the SETD3-2A protein complex in-depth we provide a striking example of how positive-stranded RNA viruses co-opt non-catalytic functions of host enzymes to complete their replication cycle.

Simultaneous mutation of three key residues in the SET interaction domain led to a complete loss of SETD3’s ability to support viral infection, even when this mutant (SET_257+266+288_) was overexpressed to levels much higher than endogenous SETD3. A double mutant in the SET interface, SET_257+266_, was partially defective in supporting viral replication. A stronger defect was perhaps expected based on the loss of 2A binding measured using affinity purification mass spectrometry. However, SET_257+266_ still retained some *in vitro* binding compared to the complete loss of binding mutant SET_257+266+288_. We speculate that this residual binding in the context of live cells is sufficient to partially rescue viral replication. Our findings that structure-guided mutations in SETD3 that ablate protein-protein interactions negate viral infection strongly suggest that SETD3’s role in viral replication is mediated solely by virtue of its interaction with the viral nonstructural protein 2A.

We have previously shown that viral entry and initial IRES translation is unaffected in SETD3 KO cells, but subsequent viral RNA genome replication is blocked^8^. Our results suggest a model in which SETD3 is a critical host factor for EV pathogenesis by promoting an early step in viral RNA replication through complex formation with the viral protease 2A. SETD3 binding to 2A may promote viral replication by regulating 2A protease activity or other molecular functions of 2A. We observed a modest (1.5-fold) increase in 2A cleavage of a P1-2A substrate peptide *in vitro* in the presence of SETD3, indicating that SETD3 is not an obligatory co-factor for 2A proteolytic activity but may enhance it. Moreover, we found that SETD3 stabilizes 2A in mammalian cells, which could affect the function of 2A in replication. It seems unlikely that the minor enhancing effect of SETD3 on 2A protease activity underlies the strong defect observed in RNA replication in SETD3 KO cells. However, we cannot exclude that SETD3 has a subtle effect on the kinetics of 2A protease activity during viral infection which could be crucial, especially early in the viral life cycle when 2A is limiting. Alternatively, SETD3 is required for another role of 2A in RNA replication. Intriguingly, the enterovirus 2A protease has been proposed to have protease-independent functions in viral RNA replication^16, 17^, although the molecular mechanisms are incompletely understood. Further work is needed to define the protease-independent functions of 2A and if SETD3 is essential for these roles.

The genetic diversity of enteroviruses, their prevalence and their ability to cause disease make them attractive for the development of HDT. SETD3 is required for EV replication in human cell lines and *Setd3* KO mice are completely protected against enterovirus pathogenesis, highlighting its promise as an antiviral target^8^. Previous studies have nominated other cellular factors as targets for HDT against enteroviruses. For example, PI4KB has been pursued because inhibitors displayed antiviral activity against a wide range of enteroviruses^34^. However, further development of PI4KB inhibitors for anti-EV therapy has been impeded by target related toxicity and mortality of mice at tested concentrations of inhibitors suggesting that even short term inhibition of PI4KB is deleterious^35, 36^. Further evidence that this toxicity was target-related came from the embryonic lethality observed in *Pik4b* knockout mice^36^, and rapid lethality in a conditional *Pik4a* knockout model upon tamoxifen administration in adult mice^37^. In stark contrast, mice with homozygous knockout alleles for *Setd3* were born without obvious defects, and developed similarly to wild type mice^10^, suggesting that inhibitors towards SETD3 may not suffer from serious toxicity issues. However, SETD3 might have more specialized functions in actin-related processes including childbirth as we observed maternal dystocia in *Setd3* knockout mice^8^. Future studies with conditional SETD3 ablation or pharmacological inhibition should be performed to assess antiviral activity, toxicity and potential pregnancy-related issues following short term inhibition of SETD3.

Our data provides insight into the molecular basis of the interaction between 2A and SETD3, and may facilitate the development of peptides or small molecule inhibitors that specifically disrupt this interaction. Chemical targeting of related protein lysine methyltransferases has largely focused on small molecules targeting the substrate-binding pocket^38^. Most protein lysine methyltransferases exhibit a high degree of substrate specificity, offering greater opportunities for selectively targeting individual enzymes. Recently, actin-based peptidomimetics were developed as inhibitors against recombinantly expressed SETD3^28^. Since the actin binding pocket and the 2A binding site partially overlap in the substrate binding region, peptidomimetic substrate inhibitors might be a starting point for the development of antivirals. We tested an actin peptide with histidine 73 replaced by leucine in an *in vitro* competition assay with 2A for binding to SETD3, and showed that actin-based peptidomimetics can inhibit the binding interaction of SETD3 with 2A. This serves as a proof of principle that actin-based peptidomimetics might inhibit viral infection. Alternatively, high throughput screening approaches could be used to discover small molecule inhibitors for the SETD3-2A interaction. Because several key residues (SET_257+266_) that were shown to be important for 2A binding did not affect actin methylation, it may be possible to identify small molecule inhibitors that disrupt the interaction with 2A while leaving the catalytic activity of SETD3 largely unaffected. We expect that the cryo-EM structure of SETD3 in complex with 2A as reported here will facilitate structure-based optimization once potential suitable SETD3 inhibitors are identified. For example, it can guide synthesis of analogues with side groups expected to disrupt 2A but not actin interactions. Finally, an alternative and promising approach that does not rely on disrupting protein-protein interactions, is to develop selective SETD3-degraders, as recently reported for another related methyltransferase^39^.

In summary, combined with the results from our previous work^8^, the atomic structure of the host methyltransferase SETD3 in complex with enterovirus 2A furthers the understanding of how enteroviruses have evolved to co-opt an essential cellular factor and provides information to facilitate the design of antivirals to combat the medically important class of enteroviruses.

## Acknowledgements

The CV-B3 infectious clone which encodes Renilla luciferase (pRLuc53CB3/T7)^40^, was a gift from F. van Kuppeveld. pCDNA3.1-CAGGS-mGFP was a gift from the Michael Lin lab. This work was supported by funds from NIH (P50 AI150476 and U19 AI135990 to N.J.K.; R01 AI140186, R01 AI141970, and R01 AI130123 to J.E.C.; T32 AI00732 to C.E.P; R35GM139569 to O.G.), the Burroughs Wellcome Fund (Investigators in the Pathogenesis of Infectious Disease to J.E.C), the National Science Foundation (GRFP Fellowship grant DGE-1656518 to C.E.P) and the Gladstone BioFulcrum Viral and Infectious Disease Research Program, and the Quantitative Bioscience Institute. We thank Peter Hwang from the Molecular Structure Group at UCSF for guidance on BLI analysis.

## Author Contributions

C.E.P., U.S.-G., K.A.V., R.H. designed, conducted, and analyzed experiments. G.M.J., J.X., and E.H.P. conducted experiments. C.E.P., U.S.-G., M.E., K.A.V., R.H., J.E.C. and N.J.K. interpreted results. M.O., O.G., K.A.V., R.H., J.E.C. and N.J.K. supervised the project. M.O., O.G., K.A.V., J.E.C. and N.J.K. obtained funding for this project. C.E.P., U.S.-G., M.E., K.A.V., R.H., J.E.C. and N.J.K. designed figures. The manuscript was written by C.E.P, U.S.-G., M.E., K.A.V., R.H., J.E.C. and N.J.K. All authors read and approved the final version of this manuscript.

## Methods

### Cells and Reagents

HEK293T/17 (CRL-11268) and H1-Hela (CRL-1958) cells were obtained from the American Type Culture Collection (ATCC). 293FT (R70007) cells were obtained from Thermo Fisher. HEK293T/17 cells were cultured in Dulbecco’s modified Eagle’s high glucose with L-Glutamine and Sodium Pyruvate (Corning) supplemented with 1× penicillin-streptomycin (Invitrogen), and 10% heat-inactivated fetal bovine serum (Invitrogen). H1-Hela and 293FT cell lines were cultured in Dulbecco’s modified Eagle’s high glucose with L-glutamine and sodium pyruvate (Gibco) supplemented with 1× penicillin-streptomycin (Sigma), and 10% heat-inactivated fetal bovine serum (Sigma).

### Affinity purification of FLAG-tagged proteins

HEK293T cells were plated 20-24 hours prior to transfection. For SETD3 affinity purifications (APs) in the presence and absence of CV-B3 2A protease, we transfected the cells with a single FLAG-tagged plenti-CMV-SETD3 construct and a 2xStrep-tagged catalytically dead CV-B3 2A pcDNA4 construct^8^. The CV-B3 2A gene sequence was derived from the CV-B3 GenBank (M33854.1) Nancy strain sequence, which annotates the starting amino acids of 2A as GQQ. For SETD3 mutant APs cells were transfected with single FLAG-tagged pLenti-CMVTRE3G-Puro-DEST SETD3 WT and mutants constructs (see below Engineering of lentiviral constructs) and a 2xStrep-tagged catalytically dead CV-B3 2A pcDNA4 construct. For each single-step purification, cells were transiently transfected using PolyJet Transfection Reagent (SignaGen Laboratories) at a 1:3 ratio of plasmid to transfection reagent. APs were performed based on previous descriptions^12, 25^. In brief, about 40 hours post-transfection cells were harvested and lysed in cold lysis buffer containing 50 mM Tris-HCl pH 7.4, 150 mM NaCl, 1 mM EDTA supplemented with 0.5% Nonidet P 40 Substitute (NP40; Fluka Analytical), complete mini EDTA-free protease and PhosSTOP phosphatase inhibitor cocktails (Roche) and incubated for 30 min. The lysate was incubated with 30 µl equilibrated anti-FLAG M2 Affinity Gel beads (Sigma-Aldrich) for 2h at 4°C. The beads were washed three times with lysis buffer containing 0.05% Nonidet P40, followed by one wash with lysis buffer without detergent. Proteins were eluted with 50 mM Tris pH 7.5, 150 mM NaCl, 1 mM EDTA containing 100 µg/ml 3xFLAG peptide (ELIM) and 0.05% RapiGest (Waters). For mass spectrometry (MS) analysis the AP eluates were reduced with 1 mM DTT at 37°C for 30 min, then alkylated with 3 mM iodoacetamide for 45 min at room temperature, followed with quenching by addition of 3 mM DTT for 15 min. The samples were digested with sequencing grade modified trypsin (Promega) overnight at 37°C. The resulting peptides were cleaned up for MS analysis using Ultra Micro Spin C18 columns (The Nest Group).

### Sequential FLAG-Strep purifications (Double affinity purifications)

Sequential purifications followed the procedure for single-step FLAG AP (described above) followed by a Strep AP. Alterations as follows: For each purification, five dishes were co-transfected and subsequently combined and lysed in proportional volumes of buffer about 40 hours post-transfection. Cell lysates were divided to perform five separate FLAG purifications. Following wash steps, FLAG beads were combined and proteins were incubated for 30 min with 0.1 mg/ml 3xFLAG peptide (ELIM) in a buffer containing Tris-HCl pH 7.4, 150 mM NaCl, 1 mM EDTA supplemented with 0.38% NP40 to elute proteins. After removing eluted protein, FLAG beads were rinsed with the same buffer for an additional elution which was combined with the initial peptide elution and incubated with MagStrep “type 3” magnetic bead suspension (IBA Lifesciences) for 2h at 4°C. After binding and washing beads, proteins were eluted using 0.1 M Glycine pH 2.5 for 10 min. Eluates were neutralized with 1 M Tris pH 9.0 and subjected to trypsin digestion followed by desalting as described above for the single-step FLAG-based AP (see above).

### Mass spectrometry analysis and scoring for global single and double AP-MS experiments

Peptides from single and double APs were resuspended in 4% formic acid and 3% ACN loaded onto a 75 μm ID column packed with 25 cm of Reprosil C18 1.9 μm, 120Å particles (Dr. Maisch). Peptides were eluted into a Q-Exactive Plus (Thermo Fisher) mass spectrometer by gradient elution at a flow rate of 300 nl/min delivered by an Easy1200 nLC system (Thermo Fisher). The gradient was run from 4.5% to 32% acetonitrile over 53 min. All MS spectra were collected with Orbitrap detection, while the 20 most abundant ions were fragmented by HCD and detected in the Orbitrap. Resulting data was searched against the SwissProt Human protein sequences (downloaded 07/2018), augmented with viral sequences. Peptide and protein identification searches, as well as label-free quantitation were performed using the MaxQuant data analysis algorithm (version 1.5.2.8 and version 1.6.12.0),^41^ and all peptide and protein identifications were filtered to a 1% false discovery rate. Variable modifications were allowed for methionine oxidation and protein N-terminus acetylation. A fixed modification was indicated for cysteine carbamidomethylation. Full trypsin specificity was required. The first search was performed with a mass accuracy of +/- 20 parts per million (ppm) and the main search was performed with a mass accuracy of +/- 4.5 ppm. A maximum of 5 modifications were allowed per peptide. A maximum of 2 missed cleavages were allowed. The maximum charge allowed was 7+. Individual peptide mass tolerances were allowed. For MS/MS matching, a mass tolerance of +/- 20 ppm was allowed and the top 12 peaks per 100 Da were analyzed. MS/MS matching was allowed for higher charge states, water and ammonia loss events. The data were filtered to obtain a peptide, protein, and site-level false discovery rate of 0.01. The minimum peptide length was 7 amino acids.

The Maxquant results from the SETD3 APs were scored for specific protein-protein interactions (PPIs) using the SAINTexpress (Significance Analysis of INTeractome) algorithm,^21, 22^ with spectral counts as the quantifying feature of the data. HEK293T cells transfected with an empty vector (not expressing any affinity tagged bait) were used as control condition, and were processed and analyzed in parallel with the bait protein expressing cells in order to avoid batch effects. Protein spectral counts for each sample were calculated as the sum from the peptide spectral counts for one protein in a given sample. To discriminate bona fide protein interactors of SETD3 in the presence and absence of CV-B3 2A, we set a Bayesian False Discovery Rate (BFDR) threshold of 0.05. To generate an overall list of candidate interactors for SETD3, we combined the proteins with an BFDR below 0.05 for both conditions (+/- CV-B3 2A expression). MSstats was used for statistical analysis to identify proteins with differential abundance in SETD3 APs in the presence and absence of CV-B3 2A expression and for SETD3 WT and SETD3 mutant APs^23^.

The data derived from the sequential APs of SETD3 and CV-B3 2A was also scored for specific PPIs using the SAINTexpress (Significance Analysis of INTeractome) algorithm^21, 22^, with spectral counts as the quantifying feature of the data as described above. HEK293T cells transfected with FLAG-tagged SETD3 and Strep-tagged EMCV 2A were used as a control condition. A BFDR threshold of 0.05 was set to identify bona fide interactors of the SETD3-2A complex. SETD3 and 2A protein abundances in the sequential affinity purifications were calculated by summing the intensities of the top three peptides per protein^42^. To estimate the stoichiometry of the SETD3 and 2A interaction, a ratio between the protein abundances was calculated.

### Protein expression and purification

A codon-optimized gene for CV-B3 (Nancy strain, M33854.1; GenBank) protease 2A (WT and C107A mutant) and for human SETD3 were separately cloned into *E. coli* expression vector 2G-T. pET His6 GST TEV LIC cloning vector (2G-T) was a gift from Scott Gradia (Addgene plasmid # 29707; http://n2t.net/addgene:29707; RRID:Addgene_29707). Single, double and triple mutants of SETD3 were created by site-directed mutagenesis using the In-Fusion cloning protocol (Takara) following manufacturer’s recommendations. Primers were designed using previously published guidelines^43^.

SETD3 and protease 2A were expressed and purified separately, and only combined for complex formation in the last purification step. Each plasmid was transformed into *E. coli* strain BL21*(DE3), then expressed overnight (300 µM IPTG) at 18°C in Luria Broth (LB) containing ampicillin. Cells were pelleted and frozen in liquid nitrogen. Purification of WT or mutant SETD3 and 2A followed the same protocol: Pellets from 3 l *E. coli* culture (2A) or 4 l *E. coli* culture (SETD3) were resuspended in lysis buffer (50 mM Tris/HCl pH 7.5, 0.2 M NaCl, 1 mM DTT) with 0.1 mg/ml lysozyme, stirred for 30 min, and lysed by sonication after addition of protease inhibitors AEBSF (GoldBio) and cOmplete EDTA-free Protease Inhibitor Cocktail (Roche). The lysate was cleared by centrifugation and the supernatant (SN) incubated for 1.5 h with Glutathione Sepharose 4B (GE Healthcare), equilibrated in buffer A (25 mM HEPES pH 7.4, 300 mM NaCl, 10% glycerol, 1 mM DTT). SN and resin were applied to a gravity column. The resin was washed with 20 column volumes (CV) of buffer A and eluted with a step gradient of buffer B (25 mM HEPES pH 7.4, 300 mM NaCl, 10% glycerol, 20 mM glutathione). Fractions containing target protein were combined. The GST-target fusion protein was digested with TEV protease for 3 h at room temperature and applied to a 5 ml His-Trap FF column (Cytiva Life Sciences). Cleaved target protein eluted in the flow-through of the column with TEV protease and GST being retained on the Ni-column.

For complex purification, separately purified 2A and SETD3 were combined at a ratio of 2:1 (mol/mol), incubated on ice for 1 h, and purified over a Superdex 200 Increase size exclusion column (Cytiva) in 20 mM HEPES pH 7.2, 150 mM NaCl, 500 µM TCEP. Aggregated material, complex of 2A with SETD3, and monomeric 2A were well separated in the column elution. Purified complex was used for structure determination by cryo-EM.

For expression and purification of WT 2A with C107A, recombinant plasmid was transformed into *E. coli* strain Rosetta(DE3)pLysS and expressed overnight (300 µM IPTG) at 18°C in Luria Broth (LB) containing ampicillin and chloramphenicol. Pellets from 4 l induced *E. coli* culture (2A) were resuspended in lysis buffer (50 mM Tris/HCl pH 7.5, 200 mM NaCl, 1 mM DTT) with 100 µg/ml lysozyme, stirred for 30 min, and lysed by sonication after addition of protease inhibitors AEBSF (GoldBio). The lysate was cleared by centrifugation and the supernatant (SN) incubated for 1.5 h with Glutathione Sepharose 4B (GE Healthcare), equilibrated in buffer A (25 mM HEPES pH 7.4, 300 mM NaCl, 10% glycerol, 1 mM DTT). SN and resin were applied to a gravity column. The resin was washed with 20 column volumes (CV) of buffer A and eluted with a step gradient of buffer B (25 mM HEPES pH 7.4, 300 mM NaCl, 10% glycerol, 20 mM glutathione). Fractions containing 2A-GST-fusion protein were combined and TEV protease added at a 25:1 ratio (w/w). The mixture was dialysed against 20 mM Tris pH 7.7, 100 mM NaCl, 1 mM DTT, 10% glycerol for 4 h, followed by anion exchange chromatography over 5 ml HiTrap Q column with buffer A (20 mM Tris/HCl pH 7.7, 1 mM DTT, 10% glycerol and buffer b (20 mM Tris/HCl pH 7.7, 1 M NaCl, 1 mM DTT, 10% glycerol). GST and 2A were well separated on the HiTrap Q column. Aliquots of purified WT 2A were frozen in liquid nitrogen and stored at −80°C.

For biolayer interferometry experiments protease 2A from CV-B3 was sub-cloned into LIC cloning vector 2Bc-T (Addgene plasmid # 37236) with a C-terminal His6-tag. pET His6 LIC cloning vector (2Bc-T) was a gift from Scott Gradia (Addgene plasmid # 37236; http://n2t.net/addgene:37236; RRID:Addgene_37236). 2A-protease in 2Bc-T plasmid was expressed as described for the 2G-T-plasmid expressed proteins. Cell lysis also followed the same protocol. Lysis SN was applied to a 5 ml His-Trap column (Cytiva) in 25 mM HEPES pH 7.4, 300 mM NaCl, 10% glycerol, 1 mM DTT, followed by a wash step with the same buffer, and gradient elution to 25 mM HEPES pH 7.4, 300 mM NaCl, 500 mM imidazole, 10% glycerol, 1 mM DTT in 20 CV. Fractions containing 2A-protease-His6 were combined and diluted 1:1 (vol/vol) with 20 mM Tris/HCl pH 8.0, 10% glycerol, 1 mM DTT). The diluted sample was purified over a 5 ml HiTrap Q column (Cytiva), equilibrated in 20 mM Tris pH 8.0, 50 mM NaCl, 10% glycerol, 1 mM DTT and eluted with a gradient over 20 CV to 20 mM Tris/HCl pH 8.0, 1 M NaCl, 10% glycerol, 1 mM DTT. Fractions with pure His-tagged 2A-protease were combined and flash frozen in liquid nitrogen until further use in biolayer interferometry experiments.

SETD3 WT and mutants used for BLI experiments were purified over glutathione-sepharose 4B, followed by TEV digest and HisTrap FF chromatography, as described in the protein purification section above. Fractions with SETD3 eluted from the HisTrap FF column were concentrated and purified by size exclusion chromatography over Superdex 200 Increase in 20 mM Tris pH 7.0, 150 mM NaCl, 5% glycerol, 1 mM DTT. SETD3 WT and various mutants showed different ratios of monomeric and aggregated SETD3 in size exclusion chromatography. Only protein from monomeric fractions was used in BLI experiments and no further concentration was performed prior to BLI experiments.

### Cryo-EM sample preparation and data collection

Purified SetD3-2A protein complex at 4.4 µM concentration in buffer consisting of 20 mM HEPES pH 7.2, 150 mM NaCl, 1 mM TCEP was applied to 300 mesh 1.2/1.3R Quantifoil grids that were previously glow discharged at 15 mA for 30 s (EMS100 glow discharger, DC negative, residual air at ∼50×10^-2^ mbar). Vitrification was done using FEI Vitrobot Mark IV (ThermoFisher) set up at 5°C, 100% humidity. 3.0 µl sample was applied on the grids and the blotting was performed with a blot force of 4 for 7 s prior to plunge freezing into liquid ethane. Cryo-EM grids were imaged using SerialEM^44^ with a 3×3 image shift collection (with calibrated correction for image shift induced beam tilt) on a Titan Krios (ThermoFisher) equipped with a K3 camera and a Bioquantum energy filter (Gatan) set to a slit width of 20 eV at nominal magnification of 105,000x with physical pixel size of 0.835 Å/pix. Camera was operated in CDS mode. Collection dose rate was 8 e^-^/pixel/second for a total dose of 66 e^-^/Å^2^ and a total exposure of 6 s and 50 ms per frame. Requested defocus range was 1.0 to 2.0 µm. Image stacks were corrected for motion and radiation damage during the data collection with MotionCor2^45^. A total of ∼1200 images were collected.

### Cryo-EM data processing

Motion corrected images were imported into cryoSPARC 2.12^46^ and CTF was estimated using patch CTF job. Micrographs were manually curated for quality and automatically picked with a circular blob of 50-100 Å. The resulting picks were curated, extracted with Fourier cropping (∼1,100,000 particles) and class averaged into 150 classes. *Ab initio* reconstruction and refinement into two classes was performed from particles contributing to 2D class averages with high resolution features (∼380,000 particles). Particles contributing to high resolution 3D class showing protein like features were re-extracted without Fourier cropping (∼260,000 particles) and refined in multiple rounds of *ab initio* reconstruction and refinement and then hetero refinement against the resulting *ab intio* reconstructions to remove low quality/contaminating particles. The final homogeneous refinement consisting of ∼100,000 particles refined to 3.6 Å and a non-uniform 3D refinement on the same particle stack yielded a 3.5 Å reconstruction. The 3.5 Å reconstruction was filtered based on the Fourier shell correlation (FSC) resolution and sharpened with a b-factor of −180 estimated based on the Guinier plot. To generate 3D FSC curves the 3DFSC server was used^47^. This reconstruction was used for downstream model building and refinement.

### Model building

Using SwissModel web-server, a homology model for 2A protease was built based on PDB:4MG3. The resulting homology model and existing SETD3 model (PDB:6MBK Chain A) were rigid body fit into the cryo-EM map in ChimeraX and then refined with phenix.real_space_refine in Phenix package^48^. The resulting model was then manually examined and corrected in ISOLDE, the SAH molecule was added into density and the model was then further relaxed into the cryo-EM density with FastRelax protocol in torsion space in Rosetta 3^49, 50^. Coot was also used for manual model examination^51^. Phenix validation was used to verify the model quality and map/model fit. Local resolution was estimated with ResMap software (ResMap was run on the two individual final half maps with the significance value of 0.05 using the automatically generated mask within cryosparc for FSC calculation)^52^. All figures were prepared in ChimeraX.^53^

### Structure analysis

SETD3 complexes were superimposed using the “align” program in PyMOL 2.3.5 (The PyMOL Molecular Graphics System PyMol Version 2.3.5 Schrödinger, LLC, New York). Intermolecular contacts were determined using the program NCONT, part of the CCP4 suite of programs^54^. Surface areas buried in complex formation were calculated with the program PISA^55^. Figures were prepared with PyMOL (The PyMOL Molecular Graphics System, Version 2.3.5 Schrödinger, LLC) and Chimera^56^.

### Biolayer interferometry

Biolayer interferometry binding data were collected on an Octet RED384 (ForteBio) and processed using the instrument’s integrated software. For SETD3 binding assays, His-tagged CV-B3 protease 2A was loaded onto Ni-NTA coated biosensors (NTA ForteBio) at 20 μg/ml in binding buffer (20 mM Tris/HCl pH 8.0, 150 mM NaCl, 5% glycerol, 1 mM DTT) for 240 s. WT and mutant SETD3 were diluted from concentrated stocks into binding buffer. After baseline measurement in the binding buffer alone, the binding kinetics were monitored by dipping the biosensors in wells containing SETD3 at the indicated concentration for 120-240 s (association step) and then dipping the sensors back into baseline/buffer for 120 s (dissociation). Data were collected for multiple concentrations of each SETD3 variant. Using the integrated software suite, we processed sample data by subtracting the negative control, aligning curves from various sample concentrations for each variant, and fitting the data using global fit with Rmax unlinked.

In order to measure the inhibitory effect of a synthetic peptide (residues 66-81, H73L, TLKYPIELGIVTNWDD) from actin on SETD3 binding to 2A-protease, we modified the standard protocol for biolayer interferometry. As before, His-tagged CV-B3 protease 2A was loaded onto Ni-NTA coated biosensors (NTA ForteBio) at 20 μg/ml in binding buffer (20 mM Tris/HCl pH 8.0, 150 mM NaCl, 5% glycerol, 1 mM DTT). Following a wash step in binding buffer, the loaded biosensors were dipped in wells containing a 3-fold dilution series of peptide in a buffer with constant SETD3 concentration. The presence of inhibitor peptide leads to a reduction in maximum binding levels of SETD3 (nm), which correlates with peptide concentration. Analysis of maximum binding against peptide concentration allows the determination of an IC50 value for inhibition of 2A binding to SETD3 by synthetic peptides. A series of binding experiments covering 40 nM to 10 μM peptide concentration at 310 nM and 620 nM SETD3 were performed. Control experiments with a scrambled peptide of the same amino acid composition (TGPLIVKWIDDYNLET) showed no inhibition effect.

### Cellular Thermal Shift Assays (CETSA)

Cellular thermal shift assays were performed as described^57^. Briefly, 293FT cells were plated 16-24 hours prior to transfection. Cells were transfected with a 2xStrep-tagged catalytically dead CV-B3 2A pcDNA4 construct^8^, and co-transfected with either a FLAG-tagged plenti-CMV-SETD3 construct or a FLAG-tagged plenti-CMV-GFP construct. Cells were transiently transfected using Lipofectamine 3000 Transfection Reagent (Thermo Fisher Scientific) at a 1:3 ratio of plasmid to transfection reagent. 48 hours post-transfection cells were harvested and resuspended in 1X PBS. Resuspended cells were subjected to thermal treatment for 3 min, lysed, and the soluble protein fraction was analyzed by Western blot.

### Western blotting analysis

Cells were lysed in 1× passive lysis buffer (Promega) and combined with 4× Laemmli sample buffer (Bio-Rad) containing 5% β-mercaptoethanol and incubated at 95 °C for 5 min. Samples were separated by SDS-PAGE on pre-cast 4–15% polyacrylamide gels (Bio-Rad) using a Bio-Rad Mini-PROTEAN gel system, and transferred onto PVDF membranes (Bio-Rad). The PVDF membranes were blocked with 5% non-fat milk dissolved in 1× PBS (Corning) containing 0.1% Tween-20 (Sigma–Aldrich). Subsequently, membranes were incubated at 4 °C overnight with a primary antibody diluted in the blocking buffer. The following primary antibodies were used to detect the presence of indicated proteins in this study: SETD3 (ab176582; 1:5,000; Abcam), Actin (A2066; 1:2,000; Sigma-Aldrich), FLAG M2 (F3165; 1:2,000; Sigma–Aldrich), H73(3-me);^10^ 1:1,000), and GAPDH (GTX627408; 1:5,000; GeneTex). After washing three times using PBS 0.1% Tween-20, membranes were incubated with secondary antibodies coupled to HRP (GTX213111-01 (anti-mouse) or GTX213110-01 (anti-rabbit); 1:10,000; GeneTex) for 1 h at room temperature. Alternatively, primary antibodies directly conjugated to HRP were used for 1 h at room temperature: Strep HRP (71591, Sigma-Aldrich), Streptactin HRP (1610381, BioRad) and FLAG M2 HRP (A8592, Sigma-Aldrich). Membranes were then washed three times before being subjected to SuperSignal West Pico PLUS Chemiluminescent Substrate or Dura Extended Duration Substrate (Thermo Fisher Scientific) peroxide solution treatment for visualization of antibody-bound proteins using a Bio-Rad ChemiDoc Touch Imaging System.

### Protease activity assay

Enzyme activity of 2A was measured using a FRET-based plate assay with a substrate peptide sequence covering the CVB3 cleavage site between P1 and 2A proteins, Dabcyl-TKTNTGAFGQQSGADY-Edans (ScenicBio, Houston TX). Peptide ends were labeled with fluorescence donor and acceptor groups Edans and Dabcyl, respectively. Proteolytic cleavage of the labeled peptide can be followed by measuring increasing fluorescence by the Edans-group, due to diminishing fluorescence quenching. Parallel proteolytic reactions were set up with mutant 2A C107A, WT 2A, WT-2A complex with WT SETD3, and 2A complex with triple mutant SETD3_257+266+288_ at 2 µ uM concentration in 40 µ ul volumes using Corning 3575 plates. Fluorescence emission at 490 nm was measured for a range of peptide substrate concentrations between 12.5 and 200 µuM in 25 mM Tris pH 7.5, 0.100 mM NaCl, 2 mM DTT, 10% glycerol, 0.2% Tween20 at 30°C over time. In addition, a standard curve of peptide completely digested by proteinase K was established to allow conversion of fluorescence into molar product concentrations. Analysis of the fluorescence data using Prism 8 (Graphpad, San Diego, CA) included (1) subtraction of fluorescence values of mutant 2A C107A from WT 2A and 2A-complex with WT SETD3 and mutant SETD3_257+266+28_, (2) linear regression to determine the slope ΔF/time, (3) conversion of ΔF/time into product conc/time, (4) non-linear regression to determine enzyme kinetics and constants Km and kcat.

### Generation of H1Hela^+CDHR^^3^ isogenic CRISPR–Cas9 SETD3 KO cell line

H1-HeLa cells were stably transduced with lentiCas9-Blast^58^ and subsequently selected using blasticidin to generate constitutively expressing Cas9 H1-HeLa cells. LentiCas9-Blast was a gift from Feng Zhang (Addgene plasmid # 52962; http://n2t.net/addgene:52962; RRID:Addgene_52962). A single Cas9-expressing H1-HeLa clone was then transduced with lentivirus without a selection marker to stably express CDHR3 C529Y (H1-HeLa^+CDHR3^). A single H1-HeLa^+CDHR3^ clone was then chosen based on RT-qPCR of CHDR3 expression and RV-C15 RNA levels for mutagenesis. The H1-HeLa^+CDHR3^ SETD3^KO^ cell line (**Fig. S4**) was engineered using the CRISPR–Cas9 strategy as previously described.^8^

### Engineering of lentiviral constructs

To generate lentiviral constructs expressing the SETD3 (Q257A + V266D) double mutant, and the SETD3 (Q257A + V266D + Y288P) triple mutant the plenti-CMV-SETD3 construct^8^ was first used as a template to incorporate the V266D mutation by Phusion site-directed mutagenesis using the following primer pairs: 5’-GTTCCCGCGATACCCTGGC-3’ and 5’-GCCAGGGTATCGCGGGAAC-3’. The SETD3 V266D mutant was introduced into EcoRV-digested pLenti-CMV-Puro-Dest (w118-1) by Gibson Assembly (New England Biolabs). The SETD3 V266D construct was then used as a template to sequentially incorporate Q257A and Y288P mutations by Phusion site-directed mutagenesis using the following primer pairs: 5’-GAGGCAAAACGCCATTCCCACAG-3’ and 5’-CTGTGGGAATGGCGTTTTGCCTC-3’ for Q257A, and 5’ CACTACTGGTCCCAACCTGGAAG-3’ and 5’-CTTCCAGGTTGGGACCAGTAGTG-3’ for Y288P. The C-terminal FLAG tag was added on in frame using the following primers: 5′-TGTGGTGGAATTCTGCAGATATGGGTAAGAAGAGTCGAGTAAAAAC-3′, and 5′-CGGCCGCCACTGTGCTGGATCTACTTATCGTCGTCATCCTTGTAATCCTCCTTAACTCCAG CAGTGC-3′. These SETD3 mutants were introduced into either EcoRV-digested pLenti-CMV-Puro-Dest (w118-1) or EcoRV-digested pLenti-CMVTRE3G-Puro-DEST (w811-1) by Gibson Assembly (New England Biolabs).

To generate lentiviral constructs expressing the SETD3 (Q257A + Y288P) double mutant the plenti-CMV-SETD3 construct^8^ was used as a template to sequentially incorporate Q257A and Y288P mutations by Phusion site-directed mutagenesis using the following primer pairs: 5’-GAGGCAAAACGCCATTCCCACAG-3’ and 5’-CTGTGGGAATGGCGTTTTGCCTC-3’ for Q257A, and 5’CACTACTGGTCCCAACCTGGAAG-3’ and 5’-CTTCCAGGTTGGGACCAGTAGTG-3’ for Y288P. The C-terminal FLAG tag was added on in frame using the following primers: 5′-TGTGGTGGAATTCTGCAGATATGGGTAAGAAGAGTCGAGTAAAAAC-3′, and 5′-CGGCCGCCACTGTGCTGGATCTACTTATCGTCGTCATCCTTGTAATCCTCCTTAACTCCAG CAGTGC-3′. This SETD3 mutant was introduced into EcoRV-digested pLenti-CMV-Puro-Dest (w118-1) by Gibson Assembly (New England Biolabs).

To generate lentiviral constructs expressing the SETD3 (G361R + F409A) double mutant, and the SETD3 (G361R +F409A + N413A) triple mutant the plenti-CMV-SETD3 construct ^8^ was first used as a template to incorporate the G361R mutation by Phusion site-directed mutagenesis using the following primer pairs: 5’-CTCGTGCCCGTATCCCCAC-3’ and 5’-GTGGGGATACGGGCACGAG-3’. The SETD3 G361R mutant was introduced into EcoRV-digested pLenti-CMV-Puro-Dest (w118-1) by Gibson Assembly (New England Biolabs). The SETD3 G361R construct was then used as a template to incorporate F409A or F409A and N413A mutations by Phusion site-directed mutagenesis using the following primer pairs: 5’- CTATTGATAGAATCGCCACCTTGGGG-3’ and 5’-CCCCAAGGTGGCGATTCTATCAATAG-3’ for F409A, and 5’-GCCACCTTGGGGGCCTCGGAATTTC-3’ and 5’-GCCCCCAAGGTGGCGATTCTATCAATAG-3’ for F409A and N413A. The C-terminal FLAG tag was added on in frame using the following primers:

5′-TGTGGTGGAATTCTGCAGATATGGGTAAGAAGAGTCGAGTAAAAAC-3′, and 5′-CGGCCGCCACTGTGCTGGATCTACTTATCGTCGTCATCCTTGTAATCCTCCTTAACTCCAG CAGTGC-3′. These SETD3 mutants were introduced to either EcoRV-digested pLenti-CMV-Puro-Dest (w118-1) or EcoRV-digested pLenti-CMVTRE3G-Puro-DEST (w811-1) by Gibson Assembly (New England Biolabs).

To generate lentiviral constructs expressing the SETD3 (Q257A+V266D+Y288P+G361R) mutant the SETD3 V266D and SETD3 G361R constructs were used as templates to incorporate Q257A and Y288P mutations by Phusion site-directed mutagenesis using the following primer pairs: 5’- GAGGCAAAACGCCATTCCCACAG-3’ and 5’-CTGTGGGAATGGCGTTTTGCCTC-3’ for Q257A, and 5’ CACTACTGGTCCCAACCTGGAAG-3’ and 5’-CTTCCAGGTTGGGACCAGTAGTG-3’ for Y288P. These SETD3 mutants were introduced to either EcoRV-digested pLenti-CMV-Puro-Dest (w118-1) or EcoRV-digested pLenti-CMVTRE3G-Puro-DEST (w811-1) by Gibson Assembly (New England Biolabs).

The lentiviral construct expressing the SETD3-5M (Catalytic Dead) (W274A/N278A/H279A/Y313A/R316M) mutant was previously constructed.^8^ To generate lentiviral constructs expressing WT SETD3 or SETD3-5M with a C-terminal FLAG tag, the following primers were used to create an in-frame fusion: 5′-TGTGGTGGAATTCTGCAGATATGGGTAAGAAGAGTCGAGTAAAAAC-3′, and 5′- CGGCCGCCACTGTGCTGGATCTACTTATCGTCGTCATCCTTGTAATCCTCCTTAACTCCAG CAGTGC -3′.

To generate lentiviral constructs expressing mGFP with a C-terminal FLAG tag, pCDNA3.1-CAGGS-mGFP (a gift from the Michael Lin lab) was used as a template and the following primers were used to create an in-frame fusion: 5′-TGTGGTGGAATTCTGCAGATATGGTGAGCAAGGGCGAG-3’ and

-3′, and 5′- CGGCCGCCACTGTGCTGGATCTACTTATCGTCGTCATCCTTGTAATCCTTGTACAGCTCGT CCATGC -3′.

### Lentiviral packaging

Lentiviral transduction was performed on designated cell lines to generate cell lines that stably express proteins of interest. Respective constructs to express proteins of interest (see ‘Engineering of lentiviral constructs’) were cloned by Gibson Assembly (New England Biolabs) into either the third-generation lentiviral Gateway destination vector (pLenti-CMV-Puro-Dest (w118-1))^59^ or pLenti-CMVTRE3G-Puro-DEST (w811-1). pLenti-CMV-Puro-DEST (w118-1) was a gift from Eric Campeau & Paul Kaufman (Addgene plasmid # 17452; http://n2t.net/addgene:17452; RRID:Addgene_17452). pLenti CMVTRE3G Puro DEST (w811-1) was a gift from Eric Campeau (Addgene plasmid # 27565). Both plasmids harbor a puromycin-resistant gene as a marker for antibiotic selection. pLenti-CMV-Puro-Dest (w118-1) harbored the high-expressing promoter, and pLenti-CMVTRE3G-Puro-DEST (w811-1) contained an inducible expression promoter that served as a low-expression promoter without the inducible transactivator present. Lentivirus was produced by co-transfection of the transgene-expressing plasmid, with a mixture of ΔVPR, VSV-G and pAdVAntage packaging plasmids into 293FT cells using TransIT-LTI (Mirus). At 48 h post-transfection, lentivirus was harvested from the supernatant and filtered through a 0.45 µm filter. Then, 1× protamine sulfate was added to the lentivirus before transducing respective cell lines overnight. H1-Hela^+CDHR3^ SETD3 KO cells that stably expressed the protein of interest were selected by treatment with 2 μg ml^−1^ puromycin (InvivoGen) along with non transduced cells as negative controls.

### Viruses

The CV-B3 infectious clone which encodes Renilla luciferase (pRLuc53CB3/T7), was a gift from F. van Kuppeveld. To propagate CV-B3-RLuc virions, the infectious clone was first linearized by MluI restriction digestion (New England Biolabs) and then purified using QIAgens QIAquick PCR purification kit. The linearized plasmid was in vitro transcribed using MEGAscriptTM T7 Transcription Kit (Invitrogen). The CV-B3-RLuc RNA was then electroporated into H1-Hela^+CDHR3^ cells using the Gene Pulser XcellTM Electroporation System. The titre of CV-B3-RLuc was determined by end-point dilution assay according to the method of Reed and Muench and expressed as 50% tissue culture infective doses (TCID_50_). Briefly, H1-Hela^+CDHR3^ cells were infected in 96 well plates and total cell lysates were harvested 3 days post infection in Renilla Luciferase Assay lysis buffer. Luciferase expression was measured using a Renilla Luciferase Assay system (E2820; Promega) to determine the presence of infection.

### Luciferase reporter virus (CV-B3-RLuc)-synchronized infection assays

Cells were seeded on 96-well plates (Greiner Bio-One) in triplicates and incubated for 16–24 h. Cells were first pre-chilled on ice for 30 min. Next, CV-B3-RLuc (MOI = 1 TCID_50_) was pre-bound to cells on ice for 60 min. The cells were then incubated at 37 °C in an incubator for 10 min to permit viral entry. Next, the cells were washed three times with PBS and cultured in complete culture media. At the indicated time points, total cell lysates were harvested in Renilla Luciferase Assay lysis buffer, and luciferase expression was measured using a Renilla Luciferase Assay system (E2820; Promega). Luciferase activity was measured by the addition of substrate, and luciferase readings were taken immediately using a GloMax 20/20 Luminometer with a 5-s integration time.

### Statistical analysis

For infection data, GraphPad Prism 8.4 (GraphPad Software) was used to perform statistical analyses. *P*-values were determined by two-way analysis of variance (ANOVA) (Holm–Sidak corrected) on log-transformed data. For MS data, statistical models implemented in MSstats were used to calculate *P*-values. In this case, *P*-values were adjusted with Benjamini Hochberg.\*\*\*\**P* ≤ 0.0001.

### Materials Availability

All unique/stable reagents generated in this study are available from the Lead Contact with a completed Materials Transfer Agreement.

### Data and Code Availability

The mass spectrometry-based proteomics data have been deposited to the ProteomeXchange Consortium via the PRIDE partner repository with the dataset identifier PXD024127 ^60^. The data can be accessed using the following reviewer account details: Username, Password: YcBvH8Hg at https://www.ebi.ac.uk/pride/login. The accession numbers for the cryo-EM structure of CV-B3 2A bound to SETD3 reported in this paper are PDB:7LMS and EMDB:23441.

## Supplemental Information Titles and Legends

**Figure S1. SETD3-2A complex stoichiometry and reconstitution.** Related to Fig. 1 and 2. **A.** Stoichiometry of SETD3-2A complex. Absolute abundances for SETD3 and 2A were estimated from the sequential affinity purification MS data by summing up the top 3 peptide intensities per protein, assuming that the top 3 peptides per protein have a similar MS response. The ratio of estimated absolute abundances was calculated for SETD3 and 2A across two biological replicates. **B.** Coomassie stained polyacrylamide gel of fractions eluted from Superdex 200 Increase size exclusion column after complex reconstitution. Molecular weight marker, column input, two fractions for peak 1 (aggregated material), four fractions for peak 2 (complex) and three fractions for peak 3 (monomeric 2A) were loaded on the gel. Lane 6 was skipped due to problems with the loading comb.

**Figure S2. Flowchart for SetD3-2A cryo-EM data collection and image processing.** Related to Fig. 2. Detailed description of data collection parameters as well as image processing.

**Figure S3. Fourier shell correlation curves and local resolution estimation for the SETD3-2A cryo-EM reconstruction.** Related to Fig. 2. **A**. Gold standard Fourier shell correlation (FSC) curves for the last round of refinement out of cryoSPARC2 indicating a 3.5 Å cryo-EM reconstruction based on 0.143 FSC cut-off. **B**. Output of the 3D FSC server plotting rotational FSCs for the reconstruction demonstrating isotropic resolution. Reported sphericity is 0.957. **C**. Cryo-EM reconstruction of SETD3-2A protein complex colored by local resolution as reported by ResMap.

**Figure S4. Example micrograph and 2D classes.** Related to Fig. 2. **A**. Example micrograph of the SETD3-2A protein complex. **B**. 2D classes during image processing.

**Figure S5. Biolayer Interferometry and FRET-based protease assay.** Related to Fig. 3. **A.** BLI Results. Binding curves for varying input concentrations (different colors indicated in legend to individual plots) for selected SETD3 variants. Binding constants as calculated by the integrated software of the Octet Red384 instrument are shown. **B.** SETD3 mutant protein purification. Elution profile from the final purification step on Superdex 200 Increase size exclusion column for SETD3 WT and mutants. Aggregated material and monomeric SETD3 peaks are indicated with arrows. **C.** Michaelis Menten analysis of 2A protease activity. Activities of 2A alone, and in complex with WT SETD3 and SETD3 triple mutant are compared. Statistical analysis using F-test to compare the curve fit under the assumption that K_m_ and k_cat_ are the same or different for 2A and 2A + SETD3 WT rejects the hypothesis that the variables are the same for the two data sets (p<0.0001).

**Figure S6. Isogenic CRISPR-Cas9 deletion of SETD3 in H1-Hela^+CDHR3^ cells.** Related to Fig. 4B and Fig. 5. Sequencing analysis of H1-Hela^+CDHR3^#1C4 SETD3^KO^#1B6 cells compared to parental H1-Hela^+CDHR3^#1C4 cells (WT). Nucleotides 289-313 of SETD3 reference sequence are displayed.

**Figure S7. SET interface mutants inhibit enterovirus infection upon overexpression.** Related to Fig. 5. Infection of WT, SETD3^KO^, or SETD3^KO^ cells complemented with SETD3 structure derived mutants under a CMV promoter. Statistics were performed on 6h timepoints. *P-*values were determined by two-way ANOVA (Holm–Sidak corrected) on log-transformed data. RLU = relative light units, n.s. = not significant, \*\**P* ≤ 0.0001

**Table S1. Mass spectrometry data of SETD3-FLAG APs.** Separate Excel File. Related to **Fig. 1A**.

**Table S2. Mass spectrometry data of SETD3-FLAG CV-B3-Strep double-APs.** Separate Excel File. Related to **Fig. 1C**.

**Table S3.** Data collection parameters and statistics on cryo-EM reconstruction and the SETD3-2A model. **Related to** Fig. 2.

**Table S4. Intermolecular contacts between SETD3 and 2A.** Related to **Figs. 2 and 3**. ^a^Contacts were determined with the program ncont in the CCP4 program suite^54^. Residues highlighted in bold were previously identified as critical for PPIs between SETD3 and 2A in a mammalian two-hybrid system^8^.

