## Supplemental figures and tables for "Structure-function analysis of enterovirus protease 2A in complex with its essential host factor SETD3"

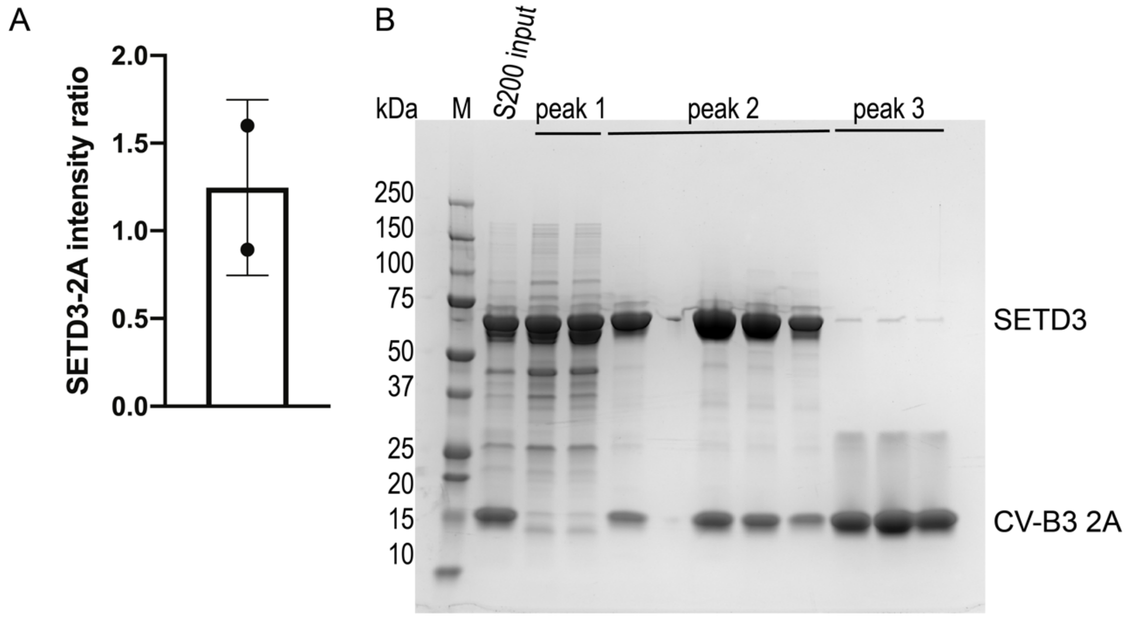

**Figure S1. SETD3-2A complex stoichiometry and reconstitution.** Related to **Fig. 1** and **2**. **A.** Stoichiometry of SETD3-2A complex. Absolute abundances for SETD3 and 2A were estimated from the sequential affinity purification MS data by summing up the top 3 peptide intensities per protein, assuming that the top 3 peptides per protein have a similar MS response. The ratio of estimated absolute abundances was calculated for SETD3 and 2A across two biological replicates. **B.** Coomassie stained polyacrylamide gel of fractions eluted from Superdex 200 Increase size exclusion column after complex reconstitution. Molecular weight marker, column input, two fractions for peak 1 (aggregated material), four fractions for peak 2 (complex) and three fractions for peak 3 (monomeric 2A) were loaded on the gel. Lane 6 was skipped due to problems with the loading comb.

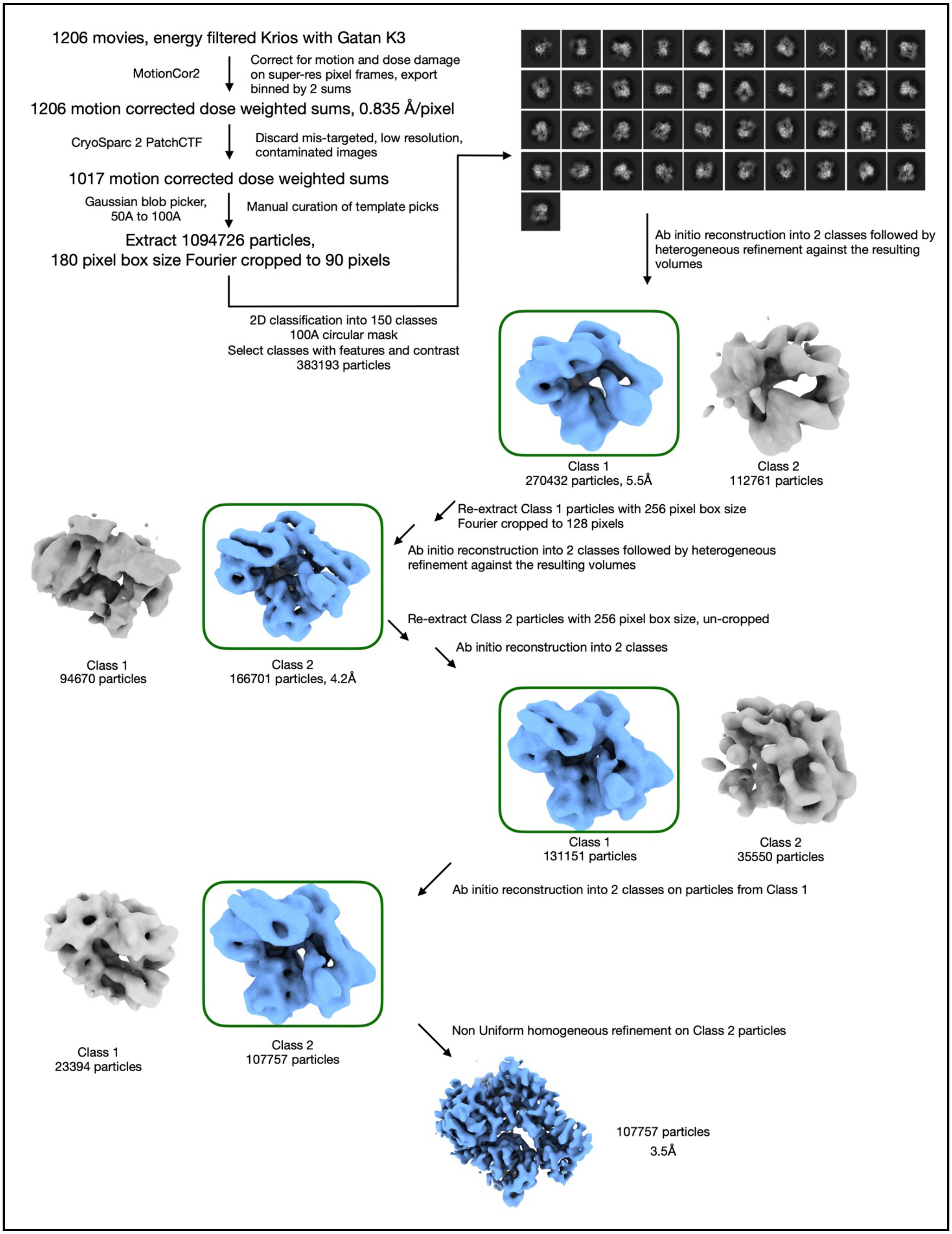
1206 movies, energy filtered Krios with Gatan K3
MotionCor2
Correct for motion and dose damage on super-res pixel frames, export binned by 2 sums
1206 motion corrected dose weighted sums, 0.835 Å/pixel
CryoSparc 2 PatchCTF
Discard mis-targeted, low resolution, contaminated images
1017 motion corrected dose weighted sums
Gaussian blob picker, 50A to 100A
Manual curation of template picks
Extract 1094726 particles,
180 pixel box size Fourier cropped to 90 pixels
2D classification into 150 classes
100A circular mask
Select classes with features and contrast
383193 particles
Ab initio reconstruction into 2 classes followed by heterogeneous refinement against the resulting volumes
Class 1
270432 particles, 5.5Å
Class 2
112761 particles
Re-extract Class 1 particles with 256 pixel box size
Fourier cropped to 128 pixels
Ab initio reconstruction into 2 classes followed by heterogeneous refinement against the resulting volumes
Class 1
94670 particles
Class 2
166701 particles, 4.2Å
Re-extract Class 2 particles with 256 pixel box size, un-cropped
Ab initio reconstruction into 2 classes
Class 1
131151 particles
Class 2
35550 particles
Ab initio reconstruction into 2 classes on particles from Class 1
Class 1
23394 particles
Class 2
107757 particles
Non Uniform homogeneous refinement on Class 2 particles
107757 particles
3.5Å

**Figure S2. Flowchart for SetD3-2A cryo-EM data collection and image processing.** Related to **Fig. 2**. Detailed description of data collection parameters as well as image processing.

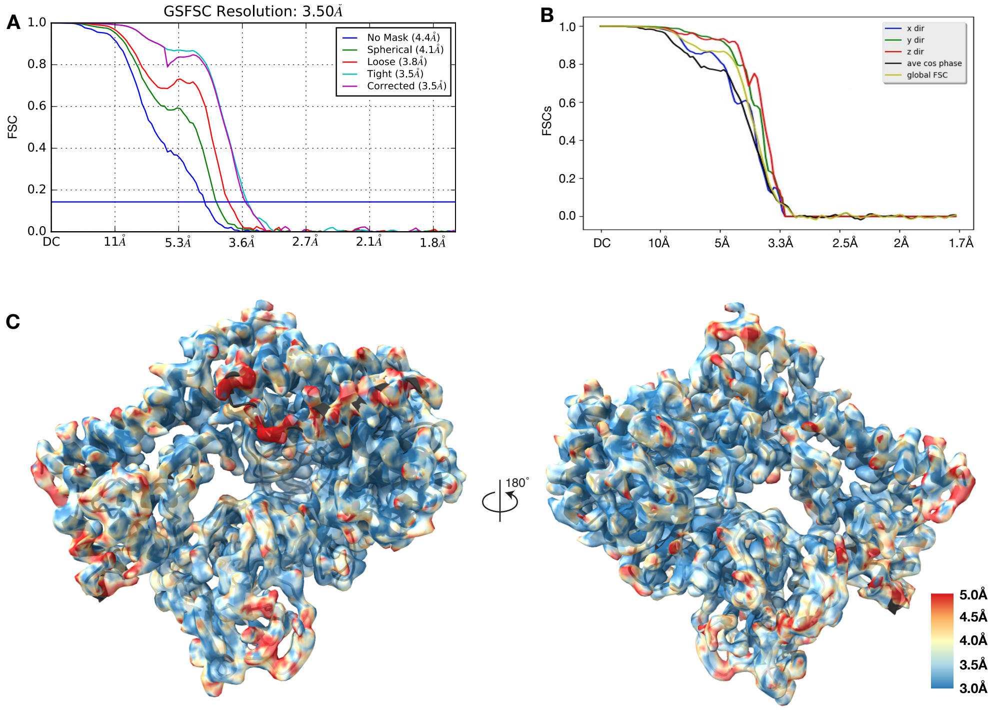

**Figure S3. Fourier shell correlation curves and local resolution estimation for the SETD3-2A cryo-EM reconstruction.** Related to **Fig. 2**. **A**. Gold standard Fourier shell correlation (FSC) curves for the last round of refinement out of cryoSPARC2 indicating a 3.5 Å cryo-EM reconstruction based on 0.143 FSC cut-off. **B**. Output of the 3D FSC server plotting rotational FSCs for the reconstruction demonstrating isotropic resolution. Reported sphericity is 0.957. **C**. Cryo-EM reconstruction of SETD3-2A protein complex colored by local resolution as reported by ResMap.

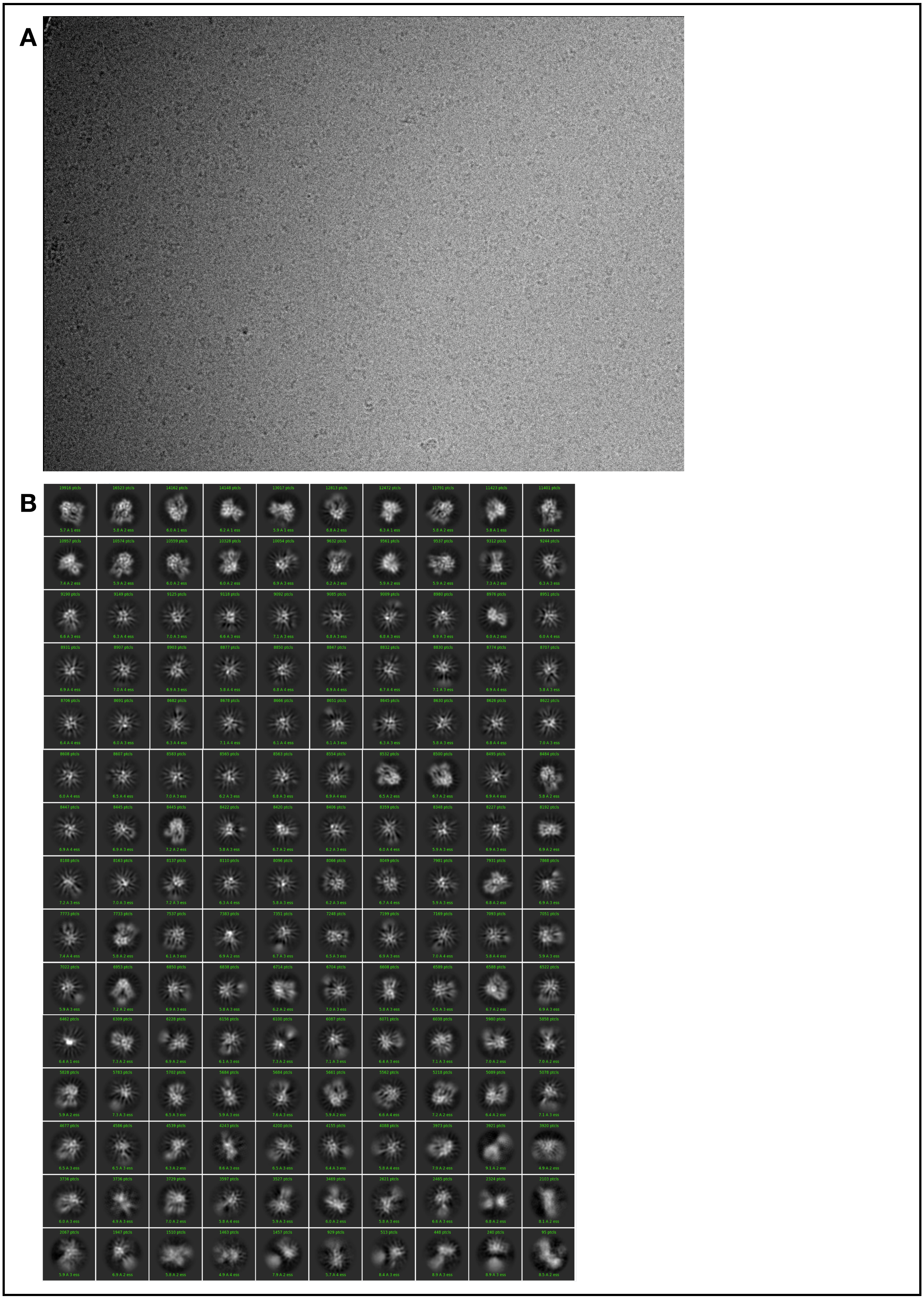
A
B

**Figure S4. Example micrograph and 2D classes.** Related to **Fig. 2**. **A**. Example micrograph of the SETD3-2A protein complex. **B**. 2D classes during image processing.

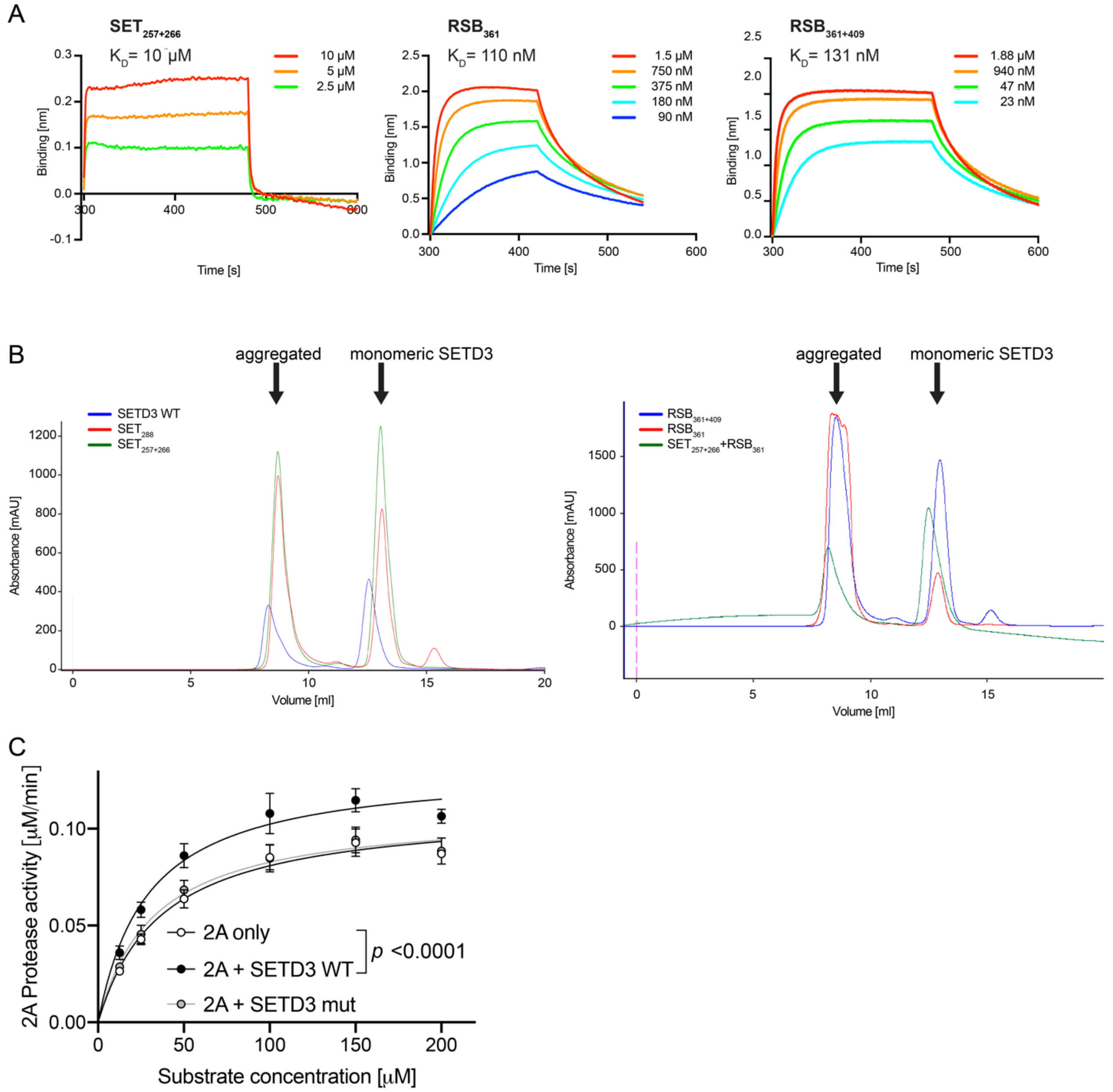

**Figure S5. Biolayer Interferometry and FRET-based protease assay.** Related to **Fig. 3**. **A.** BLI Results. Binding curves for varying input concentrations (different colors indicated in legend to individual plots) for selected SETD3 variants. Binding constants as calculated by the integrated software of the Octet Red384 instrument are shown. **B.** SETD3 mutant protein purification**.** Elution profile from the final purification step on Superdex 200 Increase size exclusion column for SETD3 WT and mutants. Aggregated material and monomeric SETD3 peaks are indicated with arrows. **C.** Michaelis Menten analysis of 2A protease activity. Activities of 2A alone, and in complex with WT SETD3 and SETD3 triple mutant are compared. Statistical analysis using F-test to compare the curve fit under the assumption that $K_m$ and $k_{cat}$ are the same or different for 2A and 2A + SETD3 WT rejects the hypothesis that the variables are the same for the two data sets ($p<0.0001$).

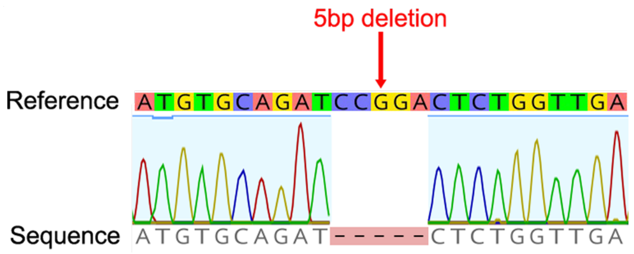

**Figure S6. Isogenic CRISPR-Cas9 deletion of SETD3 in H1-Hela$^{+CDHR3}$ cells.** Related to **Fig. 4B** and **Fig. 5**. Sequencing analysis of H1-Hela$^{+CDHR3}$#1C4 SETD3$^{KO}$#1B6 cells compared to parental H1-Hela$^{+CDHR3}$#1C4 cells (WT). Nucleotides 289-313 of SETD3 reference sequence are displayed.

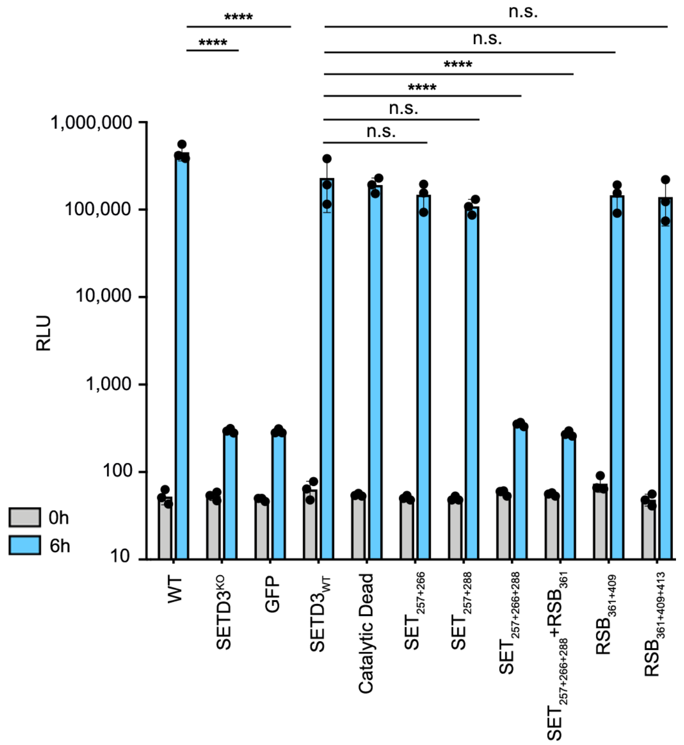

**Figure S7. SET interface mutants inhibit enterovirus infection upon overexpression.** Related to **Fig. 5**. Infection of WT, SETD3$^{KO}$, or SETD3$^{KO}$ cells complemented with SETD3 structure derived mutants under a CMV promoter. Statistics were performed on 6h timepoints. *P*- values were determined by two-way ANOVA (Holm–Sidak corrected) on log-transformed data. RLU = relative light units, n.s. = not significant, **$P \leq 0.0001$

Table S3. Data collection parameters and statistics on cryoEM reconstruction and the SETD3-2A model.

| | |
|---|---|
| **Data collection** | |
| Microscope | FEI Titan Krios |
| Voltage (keV) | 300 |
| Nominal Mag | 105,000x |
| Exposure navigation | Stage position/beam and image shift |
| Cumulative dose (e/Å^2) | 66 |
| Requested defocus range (um) | 1-2 |
| Detector | Gatan K3 |
| Detector Operation Mode | CDS |
| Pixel size (physical pixel, Å) | 0.835 |
| Dose rate ($e^-$/physical pixel/sec) | 8 |
| Total exposure time (sec) | 6 |
| Exposure per frame (sec) | 0.05 |
| Micrographs collected | 1206 |
| **Reconstruction** | EMD-23441 |
| Initial particles used | 1095000 |
| Particles selected after 2D classification | 383000 |
| Particles used in final 3D reconstruction | 108000 |
| Symmetry Imposed | C1 |
| Map Res (Å), masked/unmasked | 3.5/4.4 |
| FSC Threshold | 0.143 |
| Resolution range (local), Å | 3-5 |
| Final bfactor applied | -180 |
| **Model Refinement** | PDB-7LMS |
| Initial Model (PDB) | 4MG3, 6MBK |
| Protein residues (atoms) | 612 (4920) |
| Ligands (atoms) | 2 (27) |
| Map Correlation Coefficient (masked) | 0.83 |
| RMSD, Bond Lengths (Å) | 0.019 |
| RMSD, Bond Angles (°) | 1.576 |
| Ramachandran Outliers (%) | 0 |
| Ramachandran Allowed (%) | 0.16 |
| Ramachandran Favored (%) | 99.84 |
| MolProbity score | 0.65 |
| Clashscore (all atoms) | 0.41 |
| Rotamer outliers (%) | 0 |

Table S4. Intermolecular contacts between SETD3 and 2A.

| **SETD3 residue, # contact <4.1 Å[a]** | **2A residues** | **Hydrogen bonds/salt bridges** |
|---|---|---|
| N256, 2 | P70 | |
| Q257, 10 | **G58, V59, E114** | Q257 NE2 – E114 OE1 |
| P259, 4 | **H68** | |
| S264, 4 | **R112** | |
| R265, 5 | C47, D48 | R265 NH2 – C47 O |
| V266, 12 | **V59, R112, C113, E114** | |
| I284, 3 | F101, P70 | |
| T286, 3 | **Y69**, P70 | |
| G287, 2 | **H68** | |
| Y288, 11 | **K67, H68** | Y288 N – H68 O<br>Y288 O – H68 N |
| L290, 2 | **H68** | |
| E291, 1 | N66 | |
| E296, 3 | **K67** | E296 OE1 – K67 NZ |
| T315, 1 | S72 | |
| R336, 5 | E74 | R336 NH1 – E74 OE2<br>R336 NH2 – E74 OE2 |
| A360, 1 | S93 | |
| G361, 5 | **H94** | |
| A379, 1 | L78 | |
| Q380, 3 | P76, G77 | |
| F409, 12 | N34, L78, Y91, S93 | |
| G412, 6 | L78, Y91 | |
| N413, 10 | Y91 | |

[a] Contacts were determined with the program ncont in the CCP4 program suite (Winn et al., 2011).
Residues highlighted in bold were previously identified as critical for PPIs between SETD3 and 2A-protease in a mammalian two-hybrid system (Diep et al., 2019).
